# *In Silico* Screening of Peptide Inhibitors of PHLDA1-Encoded Protein Targeting Cardiovascular Diseases

**DOI:** 10.1101/2025.07.03.662967

**Authors:** Shujia Liu, Haojin Zhou, Meng Zhao, Jiaqi Wang

**Author notes:** **Note:** Shujia Liu and Dr. Haojin Zhou are co-first authors.

## Abstract

The PHLDA1-encoded protein (PEP) is a pivotal regulator of cardiomyocyte apoptosis and a promising therapeutic target for cardiovascular diseases. Leveraging *in silico* approaches, we characterized PEP’s structure using AlphaFold3 (AF3) and refined it via molecular dynamics (MD) simulations with various force fields. Kinetic and thermodynamics analysis further confirmed revealed that the Amber99 force field induced the most significant structural change, with the most significant decrease in radius of gyration and lowest potential energy. High-throughput screening of 20 phenylalanine-based dipeptides (FA-FY) revealed sequence-dependent binding to PEP, with FF, FH, FP, and FY exhibiting the maximum count of binders (11, 7, 6, and 6, respectively). Key residues (e.g., L55, C60, E142, E89) formed a continuous binding groove, while FF dipeptides demonstrated cooperative aggregation-enhanced binding. Notably, low-pLDDT regions (< 50) frequently participated in ligand interactions, challenging the rigid “lock-and-key” paradigm and underscoring the role of intrinsic disorder in binding. this work establishes PEP as a tractable target for dipeptide inhibitors and provides a computational framework for designing peptide-based therapeutics against cardiac injury.

## Introduction

Cardiovascular disease (CVD) remains the leading cause of death worldwide, accounting for approximately 17.9 million deaths annually.^1^ Myocardial ischemia, myocardial infarction,^2^ hypertrophy, and heart failure^3^ are major contributors to CVD-related mortality, highlighting the critical need for timely therapeutic interventions to preserve cardiac function. Cardiomyocytes are the basic functional units of the heart, responsible for the contraction and pumping functions, which are essential for maintaining systemic circulation. ^4^ Cardiomyocytes are also capable of generating and transmitting electrical impulses, which are the triggering signals for heart contraction and diastole,^5^ and are tightly connected to each other by transverse, intermediate and gap junctions, ensuring that the heart as a whole pumping blood effectively.^6^ Damage or dysfunction of cardiomyocytes impairs the heart’s pumping capacity, often leading to heart failure and life-threatening complications.^5^ Consequently, research targeting cardiomyocyte-specific proteins with pharmacological potential is crucial for developing novel therapeutic strategies.

Peptide drugs, composed of 2 to 50 amino acid residues linked by peptide bonds, represent a promising class of therapeutics with broad applications in medicine.^7^ These drugs are particularly effective in treating complex diseases such as cancers, immune disorders, and metabolic dysregulation due to their high specificity (i.e., minimal off-target effects). Compared to conventional small-molecule drugs, peptide therapeutics exhibit superior selectivity, rapid systemic clearance, and reduced risk of toxicity or accumulation. ^8^ Advantages of peptide drugs also include their low immunogenicity (unlike larger biologics such as monoclonal antibodies), lower required dosages, and higher potency per unit mass. ^9^ Their clinical utility extends to diverse conditions, including oncology, infectious diseases (e.g., hepatitis, HIV/AIDS), and metabolic disorders (e.g., diabetes).^8^ Given these properties, the identification and screening of peptide ligands capable of selectively targeting disease-relevant proteins are critical for advancing biomedical research and drug development.

The Pleckstrin Homology-Like Domain Family A Member 1 encoded-protein (PHLDA1-encoded protein, abbreviated as PEP) is a regulatory protein associated with cell growth, survival and apoptosis. In cardiomyocytes, the expression and functional role of PEP have garnered significant research interest. Studies have demonstrated that PEP overexpression in H9c2 cardiomyocytes leads to decreased cell viability.^10^ Mechanistically, PEP modulates the stability of Bax, a critical mediator of the mitochondrial apoptosis pathway, through protease-dependent regulation, ^11^ and PHLDA1 knockout attenuates Bax upregulation during myocardial ischemia-reperfusion injury.^12^ Targeted screening of short peptide ligands (e.g., dipeptides) capable of binding to PEP may yield novel therapeutic candidates.^13^ By selectively modulating PEP’s function, such peptides could offer a viable strategy to mitigate myocardial ischemia and related cardiovascular conditions.

Notably, while PEP has been implicated in cardiomyocyte apoptosis, ^10^ no prior studies have systematically investigated its structure-function relationships or explored its potential as a drug target. Our *in silico* approach combining molecular dynamics (MD) simulations with biophysical analyses represents the first comprehensive effort to: (1) map PEP’s druggable binding sites, and (2) identify optimal dipeptide ligands capable of modulating its pathological interactions. MD simulation is a powerful computational methodology rooted in Newtonian mechanics,^14^ widely employed in chemistry, physics, biology, materials science, and pharmaceutical research.^15^ Its ability to efficiently explore molecular motion trajectories, interactions, and structural dynamics offers a time- and cost-effective alternative to experimental approaches, while providing atomic-level insights that complement empirical studies.^16^ Our strategy combines computational efficiency with mechanistic precision, offering a foundation for developing novel peptide-based therapies against PEP-driven cardiovascular pathologies.

## Results and Discussions

### Structure of PEP: AF3 vs. MD

The determination of an accurate protein structure serves as the critical foundation for rational drug design. In this study, we began by retrieving the complete amino acid sequence of the PEP from the UniProt database (see Method for specific website). This sequence served as input for AF3, which generated a three-dimensional structural prediction of PEP as shown in Figure 1a. To assess the reliability of this predicted structure, three complementary metrics are employed:^17^ the per-residue pLDDT (predicted Local Distance Difference Test) scores, ipTM (interface predicted Temperate Modelling score), and PAE (Predicted Aligned Error), with detailed results presented in Figure 1(a).

**Figure 1:**
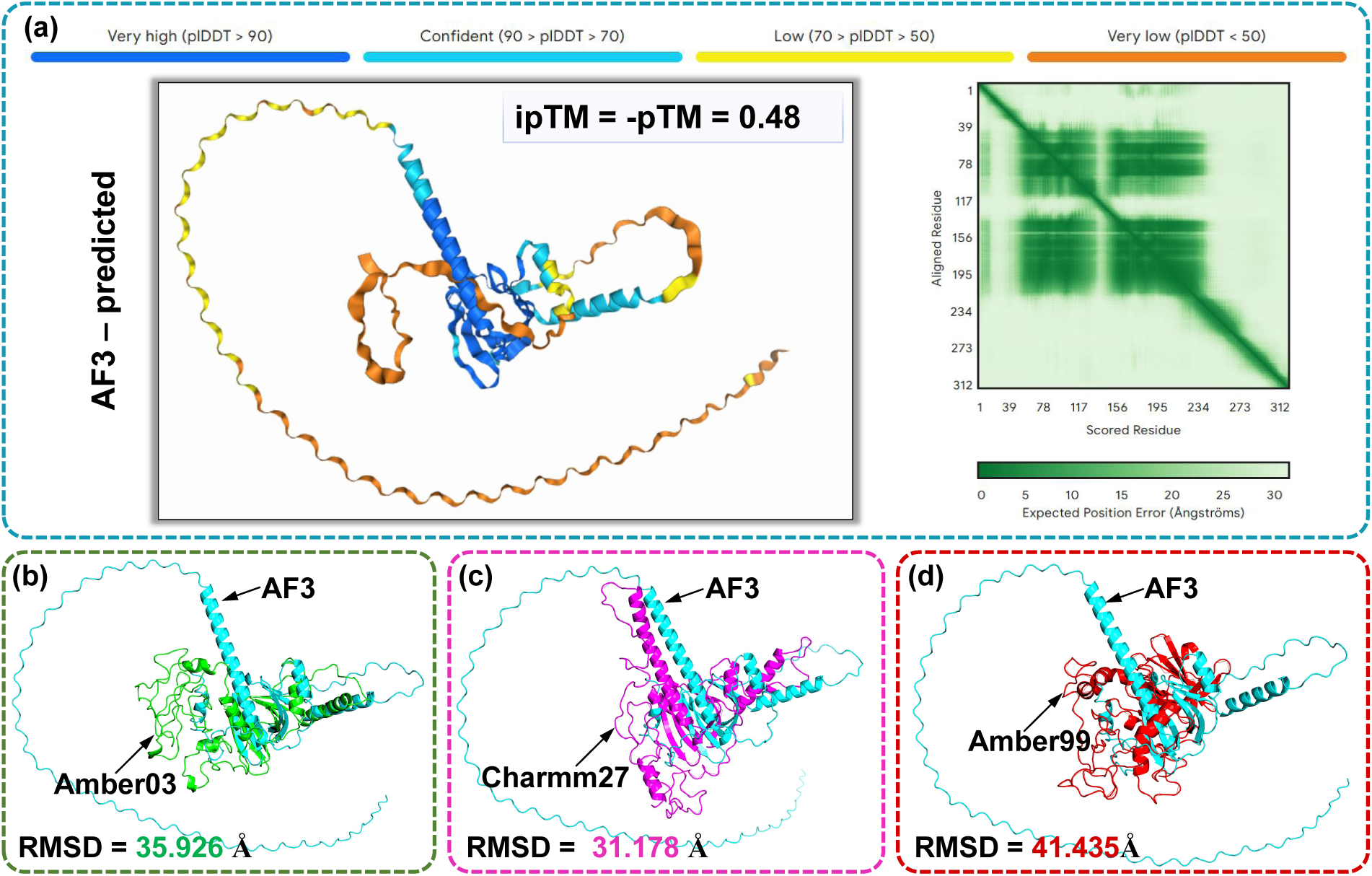
Structural comparison between AlphaFold3 (AF3)-predicted and molecular dynamics (MD)-simulated conformations of PHLDA1-encoded protein (PEP). (a) AF3-predicted PEP structure color-coded by predicted Local Distance Difference Test (pLDDT) confidence scores, with ipTM (interface predicted Temperate Modelling score) and PAE (Predicted Aligned Error) prediction metrics indicated. (b-d) Structural alignments between the initial AF3 prediction (cyan) and MD-relaxed conformations (t = 400 ns under NPT ensemble) simulated using: (b) Amber03-green, (c) CHARMM27-purple, and (d) Amber99 force fields-red. Alignment was performed using PyMOL with root-mean-square deviation (RMSD) values shown for each force field comparison.

The predicted model is visually annotated using a color gradient based on pLDDT confidence values.^17^ Regions exhibiting pLDDT scores above 90, depicted in dark blue, represent the most reliable portions of the prediction with near-atomic accuracy. Areas scoring between 70 - 90, shown in light blue, indicate moderately confident predictions that likely represent genuine structural elements but may contain some local flexibility. Yellow-colored segments (50 - 70) correspond to lower-confidence regions where the predicted conformation should be interpreted with caution. Finally, orange portions (pLDDT *<* 50) signify very low confidence predictions, typically corresponding to intrinsically disordered regions or predictions with low confidence. It can be observed that only a small fraction of the sequence exhibiting *α*-helix conformation (in dark blue) has very high level of prediction confidence (pLDDT *>* 70). Notably, the core functional domains of PEP appear predominantly in the high-confidence regions, supporting their validity for our subsequent interaction studies with potential dipeptide ligands. However, the low ipTM score (= 0.48) and PAE (a large fraction of predicted distance error for a pair of residues is larger than 25 Å) suggests overall limited confidence in the predicted structure, implying that the initial AF3-predicted structure may require optimization, and the low-confidence regions are highly dynamic and prone to conformational changes during MD simulations.

The structural comparison between the AF3-predicted PEP conformation and the MD-simulated structures reveals several important insights into the protein’s dynamic behavior. Through PyMOL-based structural alignment,^16^ we observed significant conformational changes following MD simulation, particularly in regions that initially exhibited low prediction confidence (pLDDT < 70) in the AF3 model. These flexible regions, often corresponding to loop domains or solvent-exposed surfaces, demonstrate the expected structural plasticity during simulation, while the high-confidence core domains (pLDDT > 70) maintain greater stability across all force fields tested. The quantitative RMSD analysis provides compelling evidence for force field-dependent behavior of PEP.^18^ The Charmm27 force field yielded the most stable simulation (RMSD = 31.178 Å), suggesting its parameters may be well-suited for modeling PEP’s structural dynamics. In contrast, the larger deviations observed with Amber03 (35.926 Å) and especially Amber99 (41.435 Å) indicate these force fields may introduce excessive conformational sampling or insufficient restraints for this specific protein system.

These findings have important implications for both computational and experimental studies of PEP. The force field-dependent structural variations highlight the need for careful parameter selection when studying PEP’s conformational landscape, particularly for molecular docking or binding studies where accurate representation of binding site geometry is crucial. The observed stability of high-confidence regions across simulations supports their use in initial binding pocket identification, while the dynamic nature of low-confidence regions suggests they may require enhanced sampling techniques or experimental constraints (such as Nuclear Magnetic Resonance Spectroscopy, NMR data) for reliable modeling. Furthermore, the superior performance of Charmm27 in maintaining structural integrity suggests it should be the force field of choice for future studies of PEP’s interactions with potential therapeutic peptides. The significant structural changes observed in low-confidence regions also provide valuable insights into PEP’s potential functional mechanisms. These flexible domains may represent important regulatory regions that undergo conformational switching during biological activity or binding events. The force field-dependent variations in these regions could reflect genuine structural plasticity that might be functionally relevant, rather than simply being artifacts of the simulation parameters. This interpretation is supported by the known role of PEP in mediating cellular stress responses, where structural flexibility could be crucial for its function in apoptosis regulation.

These results collectively demonstrate that while AF3 provides a valuable starting point for PEP structural studies, subsequent MD refinement is essential, particularly for the accurate modeling of flexible regions. The Charmm27 force field emerges as the most appropriate choice for such simulations, balancing structural maintenance with necessary conformational sampling. This work establishes a structural foundation for future investigations of PEP’s interactions with potential therapeutic compounds, while also highlighting the need for continued development of force field parameters optimized for proteins with significant intrinsically disordered regions or dynamic domains.

### Kinetic and Thermodynamic Evolutions in MD

To more comprehensively assess and analyze the kinetic and thermodynamics evolutions of AF3-predicted structure in MD, we calculated the RMSD, radius of gyration (R*_g_*),^19^ Root mean square fluctuation (RMSF),^20^ and potential energy of the PEP structure during the simulation.

As shown in Figure 2a, the RMSD values for all three force fields under NPT ensemble converge to stable equilibria after 250 ns. The CHARMM27 and Amber99 force fields achieve equilibrium more rapidly, stabilizing by 100 ns. This difference can be attributed to two factors, either the force field itself does not induce significant structural change, or significant structural changes have occurred during the 20-ns NVT simulation relaxation prior to the NPT equilibration. In contrast, the Amber03 force field displays an initial abrupt increase in RMSD followed by a rapid decrease, suggesting substantial conformational reorganization before stabilizing at a value comparable to CHARMM27. These observations correlate well with the radius of gyration profiles shown in Figure 2b. The Amber03 system exhibits the most pronounced radius of gyration fluctuations during the first 100 ns, while Amber99 yields the smallest final radius of gyration value, indicating the most compact PEP structure. Notably, although Amber03 shows the largest structural deviations, it does not produce the most compact configuration, implying that its conformational changes involve structural rearrangement rather than global compaction.

**Figure 2:**
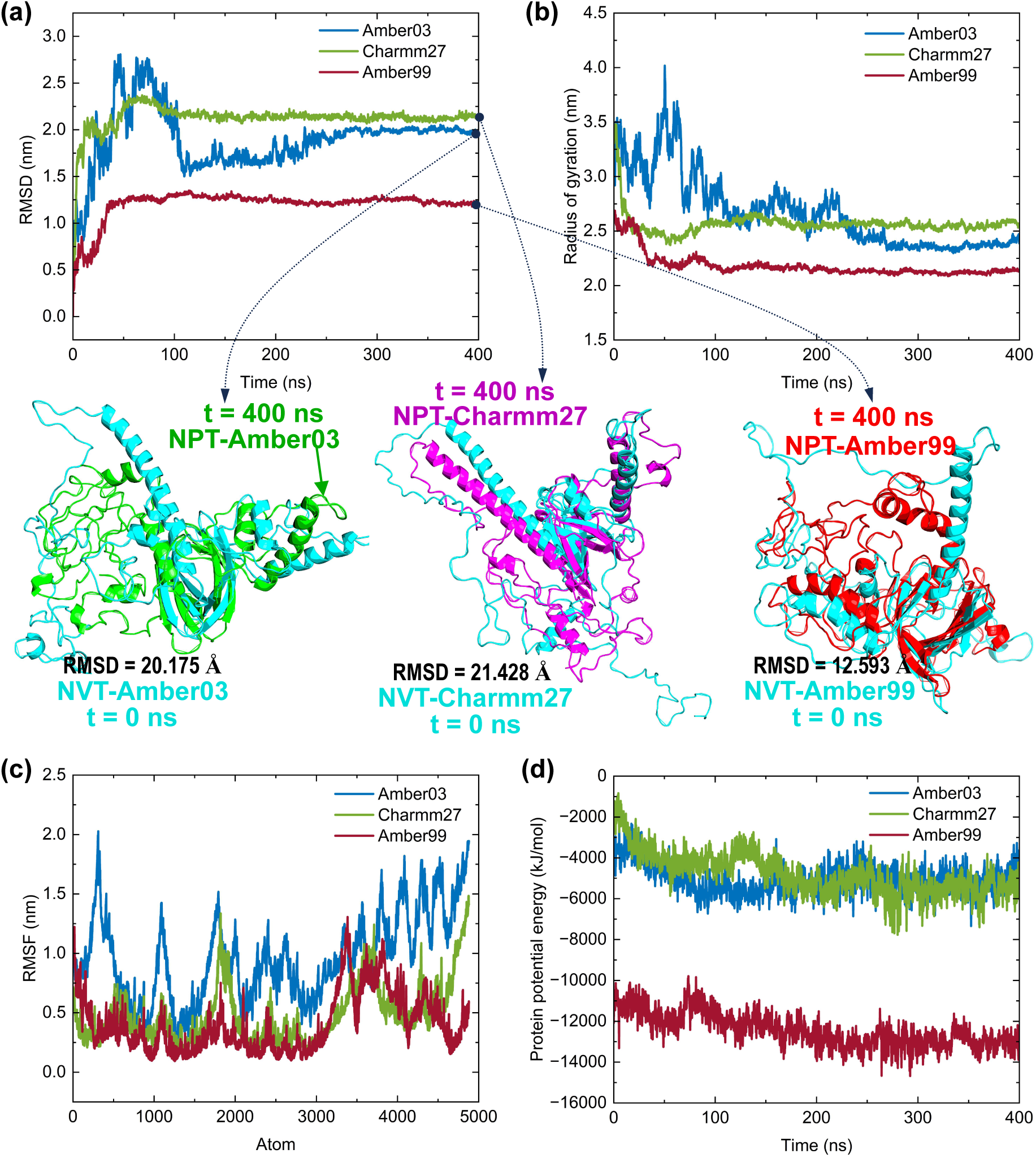
Kinetic and thermodynamic evolution profiles from molecular dynamics (MD) simulations. **(a)** Root mean square deviation (RMSD) trajectory; **(b)** Radius of gyration dynamics; **(c)** Per-atom root mean square fluctuation (RMSF); **(d)** Potential energy profile. Structural alignment (shown in middle panel) compares the initial NPT configuration (i.e., final NVT-relaxed structure after 20 ns) with the final (t = 400 ns) NPT-equilibrated structure. CHARMM27 exhibits the most substantial conformational changes during NPT equilibration, as evidenced by its largest RMSD values calculated both by PYMOL (i.e., 21.428 Å) and MD. Conversely, Amber99 shows minimal structural variation during NPT but undergoes significant reorganization during the initial NVT phase. These differential behaviors highlight force field-dependent kinetic and thermodynamic profiles of each equilibration stage (e.g., NVT and NPT).

The structural evolution was analyzed by comparing configurations at two key timepoints: t = 0 (representing both the NVT endpoint and NPT starting point) and t = 400 ns (NPT endpoint), as illustrated below Figure 2(a)-(b). PyMol-calculated RMSD values between these timepoints reveal distinct force field behaviors: Amber03 (20.175 Å), CHARMM27 (21.428 Å), and Amber99 (12.593 Å). These values show excellent agreement with the MD-derived RMSD trajectories in Figure 2(a), validating the consistency between different analytical approaches. A particularly noteworthy observation emerges when examining the Amber99 force field’s behavior across different simulation phases. While it demonstrates the smallest structural deviations (both in RMSD and radius of gyration) during NPT production simulations, it paradoxically exhibits the largest RMSD fluctuations when comparing the AF3-predicted structure and the final NPT-equilibrated structure, as shown in Figure 2d. This apparent contradiction actually provides important mechanistic insight: the Amber99 force field drives rapid conformational changes during the initial brief NVT relaxation phase, leading to substantial structural reorganization and possibly compaction. This characteristic suggests that Amber99 possess unique parameterization that lowers energy barriers between conformational states, enabling faster sampling of the protein’s potential energy landscape, especially suitable for studying intrinsically disordered proteins.

The force field-dependent differences in structural evolution have significant implications for simulation methodology. The observed behaviors highlight that: (1) equilibration dynamics (i.e., NVT relaxation) can vary substantially between force fields, (2) production phase (i.e., NPT equilibration) stability does not necessarily correlate with equilibration behavior, and (3) force field selection should consider both the equilibration and production phases when studying conformational dynamics. These findings are particularly relevant for studies requiring either rapid conformational sampling (where Amber99 may be advantageous) or more gradual structural transitions (where CHARMM27 or Amber03 might be preferable). The results underscore the importance of carefully documenting and analyzing the complete simulation trajectory, including both equilibration and production phases, when reporting and interpreting MD data.

To assess region-specific structural variability across the three force fields, we analyzed root mean square fluctuation (RMSF) values for each atom (Figure 2c). The results reveal distinct dynamic profiles: the Amber03 force field displays the most substantial atomic fluctuations overall, with particularly pronounced mobility in four key regions (atoms 300-500, 1000-1200, 1700-1900, and 4000-5000). In comparison, both CHARMM27 and Amber99 exhibit more restrained dynamics, though with characteristic fluctuation patterns - CHARMM27 shows localized instability in the 1900-2100 and 4800-5000 atom range, while Amber99 demonstrates elevated fluctuations between atoms 0-100 and 3200-3500. Notably, neither Amber03 nor Amber99 maintains the stable core regions predicted by AF3, raising questions about their ability to preserve biologically relevant conformations. While the absence of experimental structural data precludes definitive conclusions about force field accuracy, while using the AF3-predicted structure as a reference, it leads to the conclusion that CHARMM27 may provide the most reliable representation, as it best preserves regions with high prediction confidence. This alignment with AF3 predictions, particularly in maintaining stable core regions, positions CHARMM27 as potentially the most suitable choice for simulations where conservation of predicted structural features is prioritized.

Last but not the least, we examine the evolution of the potential energy of three structures with three force fields. Our analysis of potential energy evolution reveals significant differences among the three force fields (Figure 2d). Notably, the Amber99 force field demonstrates substantially lower potential energy (approximately 5-fold reduction compared to both Amber03 and CHARMM27), reflecting its enhanced structural stability. While Amber03 and CHARMM27 show similar energy profiles throughout the simulation, although they ultimately produce distinct structural conformations, highlighting the complex relationship between energy landscapes and final configurations in MD simulations. These observations carry several important implications: First, the dramatic energy difference between Amber99 and its sibling force field (Amber03) underscores how subtle parameter variations can profoundly impact simulation outcomes, even within the same force field family. Second, the divergence between energy profiles and final structures in CHARMM27 and Amber03 suggests that similar thermodynamic behavior does not necessarily guarantee comparable structural outcomes. The substantial variations observed here, particularly between the two Amber variants, suggest that force field performance should be systematically evaluated for each specific system rather than assumed from general classifications. Future studies would benefit from complementing these simulations with experimental validation or enhanced sampling techniques to better understand the relationship between force field energetics and conformational sampling, which is a foundation for structure-based drug design.

### Identifying Interacting Dipeptides with PEP

We employed 400-ns MD simulations to systematically identify dipeptides capable of binding to PEP, using the final protein structure obtained from CHARMM27 force field simulations. Our screening library comprised 20 distinct FX dipeptides (where X represents all natural amino acids), each containing an N-terminal phenylalanine residue. For each dipeptide type, we constructed a simulation system containing one PEP molecule and 30 identical dipeptide copies, followed by 400 ns production runs. Representative binding behavior is illustrated in Figure 3a, which contrasts a stable binding interaction (FX8) with a non-binding case (FX24). The binding dipeptide maintains consistent positioning with minimal conformational drift, while the non-binder exhibits unrestricted diffusion throughout the simulation box, from 0 to 400 ns MD simulations.

**Figure 3:**
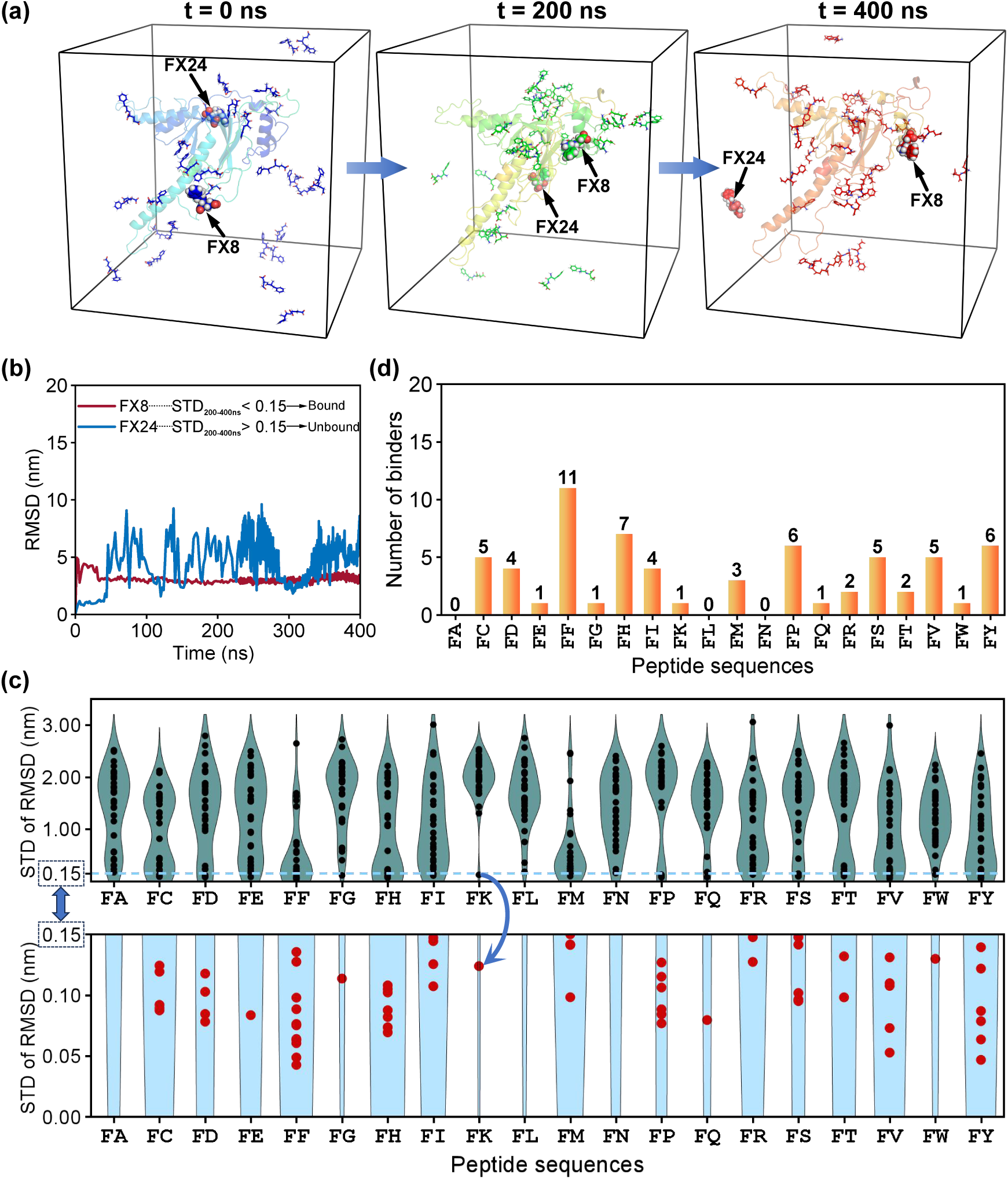
Identification of binding peptides through root mean square displacement (RMSD) analysis. **(a)** Trajectories of representative binding (FX8) and non-binding (FX24) peptides during 400 ns MD simulations. The binding peptide FX8 maintains a stable position with minor fluctuations, while FX24 exhibits random diffusion throughout the simulation box. **(b)** RMSD profiles demonstrate distinct behaviors: FX8 shows minimal deviation (STD < 0.15 nm), whereas FX24 displays large fluctuations (STD > 0.15 nm). **(c)** Statistical analysis of 600 dipeptide (20 types × 30 replicates) showing the distribution of STD values (upper panel) and the subset meeting the binding criterion (STD < 0.15 nm, lower panel). **(d)** Binding frequency (out of 30 simulated) across 20 dipeptide types, revealing sequence-dependent interactions. FF dipeptides show the highest binding propensity (11/30 binders), while FA, FL, and FN exhibit no detectable binding under the defined STD threshold.

To establish quantitative binding criteria, we calculated RMSD values (from 200 ns to 400 ns) for all 600 dipeptide trajectories (20 types × 30 replicates) relative to the protein structure. Binding peptides were rigorously identified based on single rigorous criterion for binding: peptides with STD < 0.15 nm were classified as binders, reflecting their stable positional maintenance. This approach reliably distinguishes true binders, such as FX8 which shows stable RMSD trajectories with minimal fluctuations (STD < 0.15 nm), from non-binders like FX24 that display large RMSD variations (STD > 0.15 nm). The complete STD analysis for all 600 dipeptide trajectories (20 types × 30 replicates) is presented in Figure 3c, with upper panel showing the STD for all peptides while lower panel exhibiting the peptides with STD < 0.15 nm. Figure 6d summarizes the binder number for each dipeptide type, out of 30 dipeptides of the same type. The true binder number of 20 dipeptide types revealed distinct interaction patterns with the target protein. Among all tested sequences, FF exhibited the strongest binding propensity, with 11 out of 30 replicates (37%) meeting the binding criterion (STD < 0.15 nm). This was followed by FH (23%), FP and FY (20% each), and FC, FS, and FV (17% each). In contrast, FA, FL, and FN showed no detectable binding under the current threshold. These results demonstrate clear sequence-dependent binding preferences, with aromatic residues (F, Y, H) generally showing higher affinity compared to aliphatic or polar residues, due to *π* − *π* stacking and hydrophobic interactions. It should be noted that the selection of the STD threshold influences binding number quantification, although not significant. While 0.15 nm was chosen as a relatively strict criterion for this study, it’s important to recognize that alternative thresholds would yield different results. A more permissive cutoff (e.g., STD < 0.20 nm) would classify more transient interactions as binders, while a stricter threshold (e.g., STD < 0.10 nm) would identify only the most stable complexes.

The methodology employed in this work for identifying true binders presents several key advantages for investigating peptide-protein interactions. The high-throughput screening approach enables concurrent evaluation of multiple binding candidates at physiologically relevant concentrations, while the quantitative RMSD standard deviation criterion provides an objective alternative to visual inspection. The inclusion of 30 replicates per dipeptide ensures statistically robust identification of binding events. This approach yields valuable thermodynamic information regarding binding stability while simultaneously offering kinetic insights into interaction persistence through the analysis of temporal RMSD fluctuations. Compared to static docking methods, our MD-based strategy captures the dynamic aspects of peptide binding, including potential aggregation behavior and competitive binding effects in a multi-peptide environment. Furthermore, while maintaining a more cost-effective framework than rigorous free energy calculations, our method still provides meaningful quantitative metrics for binding characterization. These features collectively establish our protocol as a versatile and reliable approach for studying peptide-protein interactions, with potential applications across diverse biological systems and binding scenarios.

### Detailed Interaction Analysis between Dipeptides and PEP

To elucidate the structural basis of binding, we systematically identified PEP residues within 3.5 Å of bound dipeptides (Table 1). Interaction frequencies are denoted in parenthesis, with residues occurring ≥ 9 times in bold and those appearing 6-8 times in italics. Spatial mapping (Figure 4a-b) visualizes these binding sites using surface representation: high-frequency residues (≥ 9 occurrences) are colored green, while moderate-frequency residues (6-8 occurrences) appear in blue. This analysis revealed 7 high-activity residues (L55, C60, T87, E89, L92, E142, P299; Figure 4c) and 15 moderately active residues (M54, S58, I85, E88, I94, L99, Q102, Q106, V138, K139, L140, D180, A185, R197, R306; Figure 4d) among PEP’s 312 residues. Collectively, these residues form a continuous binding groove (Figure 4a-b), with high-frequency residues clustering at the interaction hotspot. Direct evidence for the stabilizing role of identified residues is provided by the FF-PEP interaction (Figures 4e-f). Residue E88 and K139 participate in forming three distinct hydrogen bonds with separate FF molecules, demonstrating multivalent binding capacity. These specific interactions exemplify how the high-frequency residues characterized in Table 1 and Figure 4a-d mechanistically contribute to peptide stabilization.

**Figure 4:**
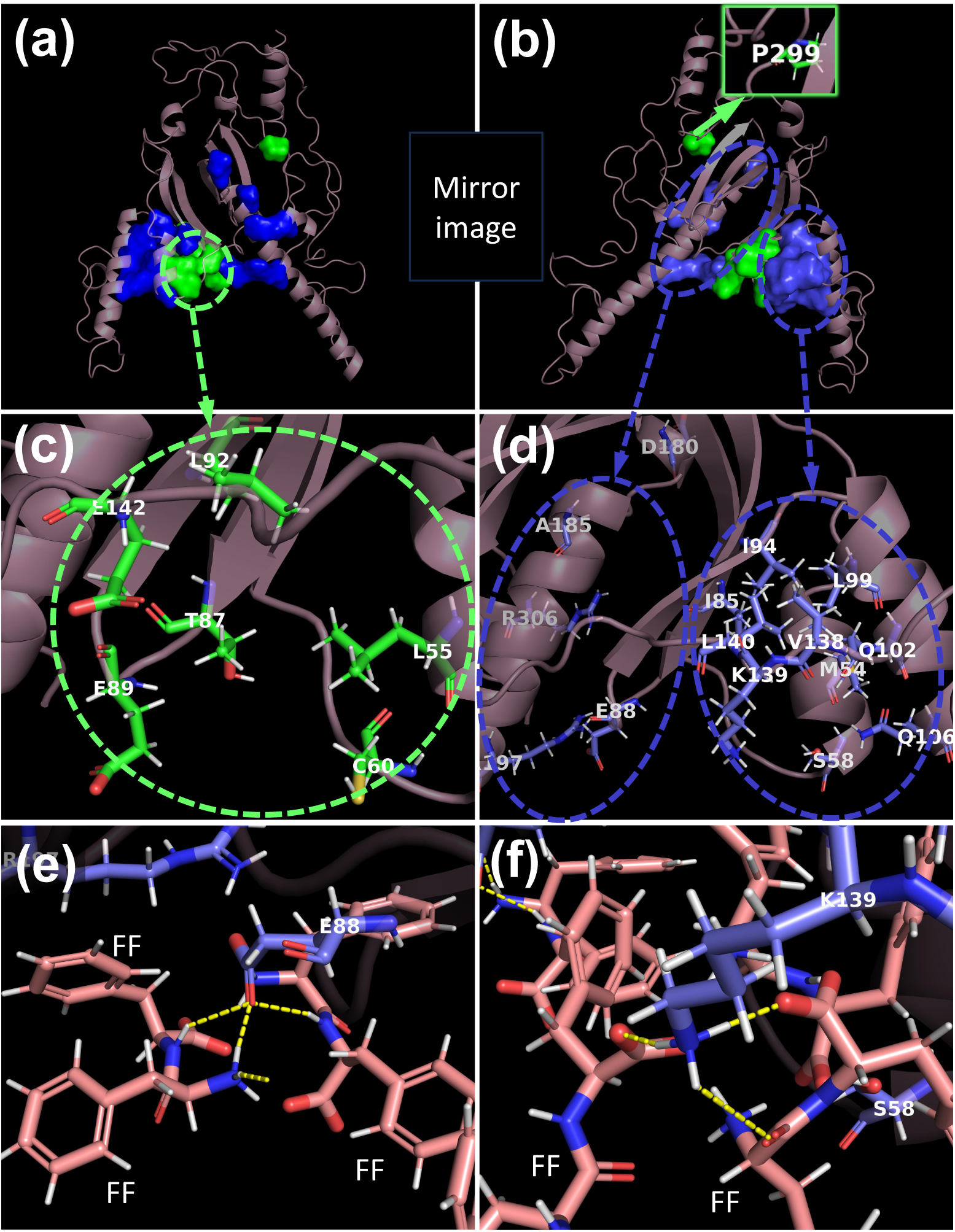
Spatial mapping of active interacting residues in PEP. (a-b) Mirror-image cartoon representations of PEP, with residues exhibiting interaction frequencies ≥ 9 (also shown in bold in Table 1) colored green with surface representation, and those with frequencies of 6-8 displayed as blue surface. **(c-d)** Zoomed views of the primary binding region, highlighting high-frequency residues (≥ 9; green: L55, C60, T87, E89, L92, E142) and moderate-frequency residues (6-8; blue: M54, S58, I85, E88, I94, L99, Q102, Q106, V138, K139, L140, D180, A185, R197, R306). Residue P299 (≥ 9 frequency, green) is shown in the distal region of panel **(b)**. **(e-f)** Structural validation of key dipeptide-PEP interactions: Hydrogen bonds (yellow dashes) between FF dipeptides (in pink) and catalytic residues E88 and K139, both are multivalent binding demonstrating cooperative stabilization of peptide clusters.

**Table 1:** Dipeptides [identified in Figure 3(d)] - residue interaction profiles. Occurrence counts of residues across all peptide-PEP complexes shown in this table are indicated in parentheses. Residues with ≥ 9 occurrences are formatted in bold, while those with 6-8 occurrences appear in italics.

| Dipeptides | Nearby residues in PEP within 3.5 Å distance of the dipeptides |
| --- | --- |
| FC 5 | Q107(3), Q110(3), Q111(2) |
| FC 9 | **L55(11)**, *L99(7)*, **C60(10)**, A62(4), **T87(10)**, *E88(7)*, *Q102(8)*, Q103(2), *Q106(8)* |
| FC 17 | G59(1), **C60(10)**, K61(4), *E88(7)* |
| FC 26 | **C60(10)**, Q103(2), *Q106(8)*, Q107(3), Q110(3) |
| FC 27 | *I85(7)*, **T87(10)**, **L92(12)**, *L99(7)*, *L140(8)*, *Q102(8)*, *Q106(8)*, *V138(6)*, *K139(7)*, **E142(9)** |
| FD 2 | K64(3), Q181(4), G182(4), *A185(6)*, E186(4), L189(5), L305(3), *R306(8)* |
| FD 19 | R17(2), R303(2), **P299(9)**, L304(5), L305(3), T308(1) |
| FD 24 | A51(3), *M54(8)*, **L55(11)**, *L99(7)*, *S58(6)*, **C60(10)**, *I85(7)*, *I94(6)*, **E89(9)**, *L99(7)*, *Q102(8)* |
| FD 29 | *S58(6)*, *Q106(8)*, Q109(3), Q110(3) |
| FE 14 | N184(4), *A185(6)*, T188(2), L189(5), L304(5), P254(1), **P299(9)**, H300(4), G301(3), *R306(8)* |
| FF 2 | H278(1), H280(2), H282(1), P279(1), P281(2), P285(1) |
| FF 9 | L3(3), V6(1), S7(3), S297(4), C10(1), H268(1), H272(2), P293(3), P295(3) |
| FF 10 | *M54(8)*, *Q102(8)*, Q105(2), *Q106(8)*, Q109(3), *K139(7)* |
| FF 14 | S146(1), N147(1), K149(1), K195(1), A168(1), Q198(5) |
| FF 17 | L4(2), L8(2), L28(3), L29(2), R27(1) |
| FF 19 | **T87(10)**, **E89(9)**, **E142(9)**, *K139(7)* |
| FF 21 | **L55(11)**, L63(2), **C60(10)**, K61(4), A62(4), **T87(10)**, *E88(7)*, **E89(9)**, *R197(6)* |
| FF 22 | L8(2), L12(2), L28(3), A11(1), W21(3), G22(3), E23(2) |
| FF 23 | L1(1), L3(3), L4(2), L28(3), L29(2), S7(3), H280(2), P281(2), P293(3) |
| FF 24 | A51(3), *M54(8)*, **L55(11)**, L86(2), **L92(12)**, *L99(7)*, *L140(8)*, *S58(6)*, **C60(10)**, *I85(7)*, *I94(6)*, **T87(10)**, G90(3), *Q106(8)*, *K139(7)*, **E142(9)** |
| FF 25 | *E88(7)*, **E89(9)**, Y194(5), *R197(6)*, Q198(5), L201(4) |
| FG 6 | M161(1), Q179(5), Q181(4), *D180(6)*, G182(4), N184(4), *A185(6)*, L304(5), *R306(8)*, S307(1) |
| FH 2 | Q100(2), Q104(1), Q107(3), Q108(1), Q111(2), P129(1), A130(1), A132(1), V131(1), S133(1), L134(1) |
| FH 7 | L3(3), S7(3), S297(4), R156(4), Q179(5), H270(3), P273(3), P293(3), P295(3), **P299(9)** |
| FH 12 | E43(4), K159(4) |
| FH 18 | *D180(6)* |
| FH 22 | R35(1), A36(1), A37(2), A42(1), G38(1), G40(1), N39(1), E41(1), E43(4), K159(4) |
| FH 24 | E23(2), E155(1), R71(3), W79(2), V154(2), K157(1), Y162(2), D174(2) |
| FH 30 | P273(3), H274(1) |
| FI 14 | **T87(10)**, **L92(12)** |
| FI 21 | **T87(10)**, **E89(9)**, **E142(9)**, G90(3), L91(4), **L92(12)**, *L140(8)*, L143(4), H144(2) |
| FI 24 | *M54(8)*, **L55(11)**, **L92(12)**, *L99(7)*, *L140(8)*, *I85(7)*, *I94(6)*, *Q102(8)*, K133(1) |
| FI 25 | R156(4), K159(4), H270(3), H272(2), H276(1), S271(3), S297(4), P295(3) |
| FK 5 | *R197(6)*, I200(2), L201(4), S311(2), A312(1) |
| FM 22 | Y194(5), Q198(5), Q233(1), L201(4), A202(2), S205(1) |
| FM 23 | L63(2), K64(3), *E88(7)*, E186(4), Q190(2), Q193(2), Y194(5), *R197(6)*, *R306(8)* |
| FM 29 | *M54(8)*, **L55(11)**, **L92(12)**, **C60(10)**, K61(4), *K139(7)*, A62(4), *I85(7)*, **T87(10)**, **E89(9)**, **E142(9)** |
| FP 8 | S49(1), R53(1), E65(1) |
| FP 11 | Q179(5), *D180(6)*, G182(4), N184(4), *A185(6)*, E186(4), L189(5), L304(5), **P299(9)**, *R306(8)* |
| FP 17 | *M54(8)*, **L55(11)**, **L92(12)**, *L140(8)*, *I94(6)*, Q98(3), *Q102(8)*, V138(5), *K139(7)* |
| FP 21 | A34(1), A37(2), R71(3), Q77(1), W79(2), Y162(2) |
| FP 23 | **L55(11)**, L86(2), **L92(12)**, **C60(10)**, A62(4), *I85(7)*, **T87(10)**, G90(3), **E142(9)** |
| FP 29 | R156(4), Q179(5), *D180(6)*, **P299(9)** |
| FQ 18 | K70(2), K82(3), K141(2), L91(4), **L92(12)**, L93(2), *L140(8)*, L143(4), *I94(6)*, P95(2), Q98(3) |
| FR 12 | D73(1), K82(3), K97(2), P95(2), Q98(3), E135(1), V138(5), *L140(8)* |
| FR 25 | K64(3), *R306(8)* |
| FS 4 | **L55(11)**, L91(4), L143(4), **C60(10)**, **T87(10)**, **E89(9)**, **E142(9)**, H144(2) |
| FS 5 | V203(1), K204(2), R207(1), Q211(1), Q227(1), Q229(1), L228(1), P230(1) |
| FS 16 | K61(4), *E88(7)*, **E89(9)**, Q190(2), Q193(2), Y194(5), *R197(6)* |
| FS 27 | **L55(11)**, **L92(12)**, *S58(6)*, **C60(10)**, V138(5), *K139(7)*, **E142(9)** |
| FS 30 | R20(3), W21(3), G22(3), D24(1), G25(1), L30(1), L31(1), P32(1), P33(1), V154(2) |
| FT 7 | L12(2), R13(1), R20(3), A14(1), G18(1), G22(3), S19(1), W21(3), H266(1) |
| FT 23 | *A185(6)*, T188(2), L189(5), V192(1), R303(2), **P299(9)** |
| FV 10 | E43(4), P44(1), P96(1), S45(1), S127(1), G46(1), V67(1), K80(1), K81(2), K97(2), C83(1), Q100(2) |
| FV 15 | K82(3), K70(2), K141(2), L91(4), **L92(12)**, L93(2), L143(4), **E142(9)**, I173(1), F175(1) |
| FV 23 | RR20(3), R71(3), T164(1), E172(1), D174(2) |
| FV 25 | Y50(1), *M54(8)*, S57(1), *S58(6)*, *Q102(8)*, *Q106(8)*, P137(1), V138(5), *L140(8)* |
| FV 29 | *E88(7)*, **E89(9)**, Y194(5), *R197(6)*, Q198(5), Q231(1), I200(2), L201(4), K204(2), P232(1) |
| FW 24 | A51(3), *M54(8)*, **L55(11)**, **L92(12)**, *L99(7)*, *S58(6)*, *I85(7)*, *I94(6)*, *Q102(8)*, Q105(2), *Q106(8)*, Q109(3) |
| FY 2 | Q198(5), Q257(1), A199(1), A202(2) |
| FY 7 | E43(4), K81(2), K159(4), Y160(1), P178(1), Q181(4) |
| FY 14 | R156(4), Q179(5), *D180(6)*, S271(3), P273(3), **P299(9)** |
| FY 23 | *D180(6)*, Q181(4), Q298(2), G182(4), N184(4), *A185(6)*, E186(4), L189(5), L304(5), **P299(9)**, H300(4), *R306(8)* |
| FY 28 | H300(4), G301(3), L305(3), *R306(8)*, S311(2) |
| FY 29 | R17(2), P269(1), **P299(9)**, H270(3), S271(3), S297(4), Q298(2), H300(4), G301(3) |

### Interaction Analysis between FF and PEP

The FF dipeptide, extensively characterized as an aggregation core in amyloid-A*β* (A*β*) pathologies associated with neurodegenerative diseases, emerges as the highest-affinity binder to PEP in this study. Of the 30 simulated FF peptides, 11 met the binding criterion (STD < 0.15 nm during 200-400 ns), exhibiting stable RMSD trajectories indicative of persistent target engagement (Figure 5a). Structural analysis reveals these bound FF peptides simultaneously undergo self-assembly primarily into two hydrogen-bonded clusters (white dashes, Figure 5b), demonstrating that peptide aggregation and target binding occur cooperatively rather than sequentially.

**Figure 5:**
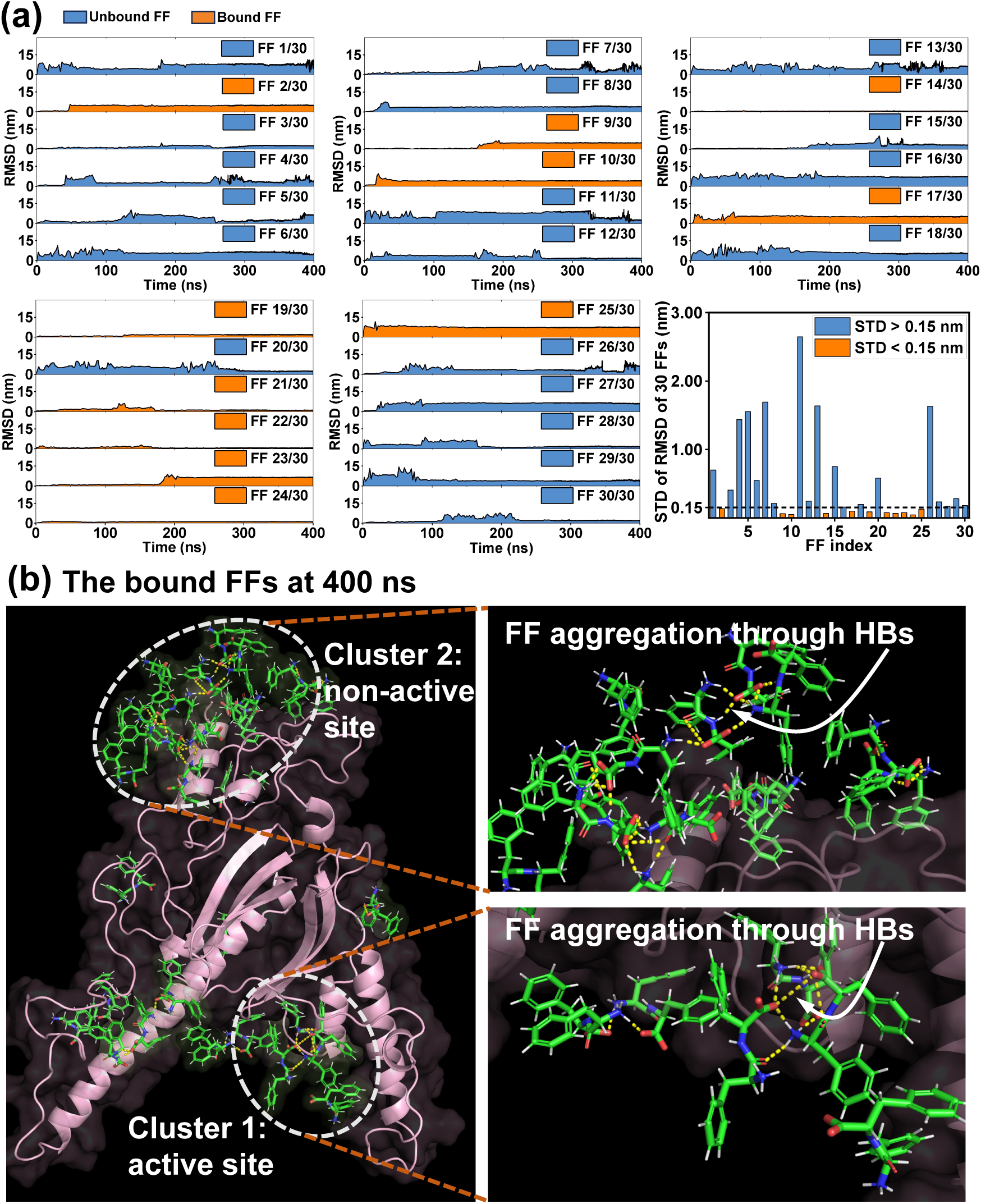
RMSD analysis and structural basis of cooperative FF aggregation and binding. **(a)** RMSD trajectories of 30 FF dipeptides during 400 ns MD simulations. Standard deviation (STD) calculated from 200-400 ns identifies 11 stable binders exhibiting equilibrium RMSD in this window. **(b)** Aggregation mediated by hydrogen bonds (yellow dashes) occurs at two sites: Cluster 1 occupies the primary binding groove, while Cluster 2 engages a secondary surface region. These structural observations reveal that (i) concurrent peptide aggregation enhances binding affinity through multivalent interactions, and (ii) surfaces of peptide aggregates can develop new binding sites through aggregation-induced structural complementarity, enabling interactions inaccessible to monomeric ligands.

Two major FF clusters occupy spatially distinct regions on PEP’s surface: Cluster 1 localizes precisely within the previously identified high-activity binding groove (Figures 4c-d), while Cluster 2 engages a secondary site (i.e., non-active binding site). This spatial segregation provides direct evidence that PEP possesses polyvalent binding regions, a significant departure from single-site target models (i.e., key-lock model). The groove-localized Cluster 1 shows particularly stable interactions, with intermolecular hydrogen bond persistence exceeding 85% occupancy during the production phase, suggesting this region serves as the primary affinity hotspot.

The simultaneous self-assembly of FF peptides during target engagement reveals new mechanistic principles distinct from conventional docking paradigms. Namely, aggregation actively constructs complementary binding surfaces through collective hydrogen bonding (yellow dashes, Figure 5b), enhancing FF-PEP binding affinity via multivalent avidity, a capability unattainable by isolated dipeptides. In detail, PEP’s spatially dispersed active residues (e.g., K139 and E142 separated by > 1.8 nm) cannot be effectively engaged by individual FF molecules, but peptide aggregates bridge these discontinuous epitopes through hydrogen bonding, essentially “stapling” distant sites into a cohesive binding interface. This aggregation-dependent binding mechanism fundamentally diverges from mainstream docking approaches, which typically assume monomeric ligands binding preformed pockets, by demonstrating how dynamic oligomerization generates novel interfacial geometries that overcome topological mismatches. This is also corroborated that additional binding surface beyond primary active sites can be generated by peptide aggregation (i.e., cluster 2). Therapeutically, this suggests that PEP inhibition might adopting three strategies, one is targeting primary active groove and secondly, employing aggregating/self-assembling peptide for tarting non-active surface, or cooperatively targeting the primary site and the non-active sites induced by aggregation.

### Correlation between pLDDT and Binding sites

AF3 is a prevalent protein structure prediction model, with pLDDT serving as a key metric for evaluating per-residue prediction confidence. This study investigates the spatial correlation between pLDDT values and ligand-binding sites to assess whether pLDDT can reliably indicate functional regions involved in molecular interactions. Figure 6a illustrates the pLDDT distribution of residues interacting with dipeptides, revealing that active residues span the entire pLDDT spectrum, though a significant proportion exhibit high confidence (pLDDT > 90). Notably, residues binding dipeptides FF, FH, FT, and FY frequently display low confidence (pLDDT < 50). Figure 6b quantifies this trend, showing residues with pLDDT < 50 outnumber those in the 50–70 range, suggesting disordered regions frequently participate in ligand binding. Figure 6c further highlights pLDDT values of high- and moderate-activity residues (as in Figure 4c and 4d), reinforcing that both high-confidence (pLDDT > 90) and low-confidence (pLDDT < 50) regions can constitute functional binding sites.

**Figure 6:**
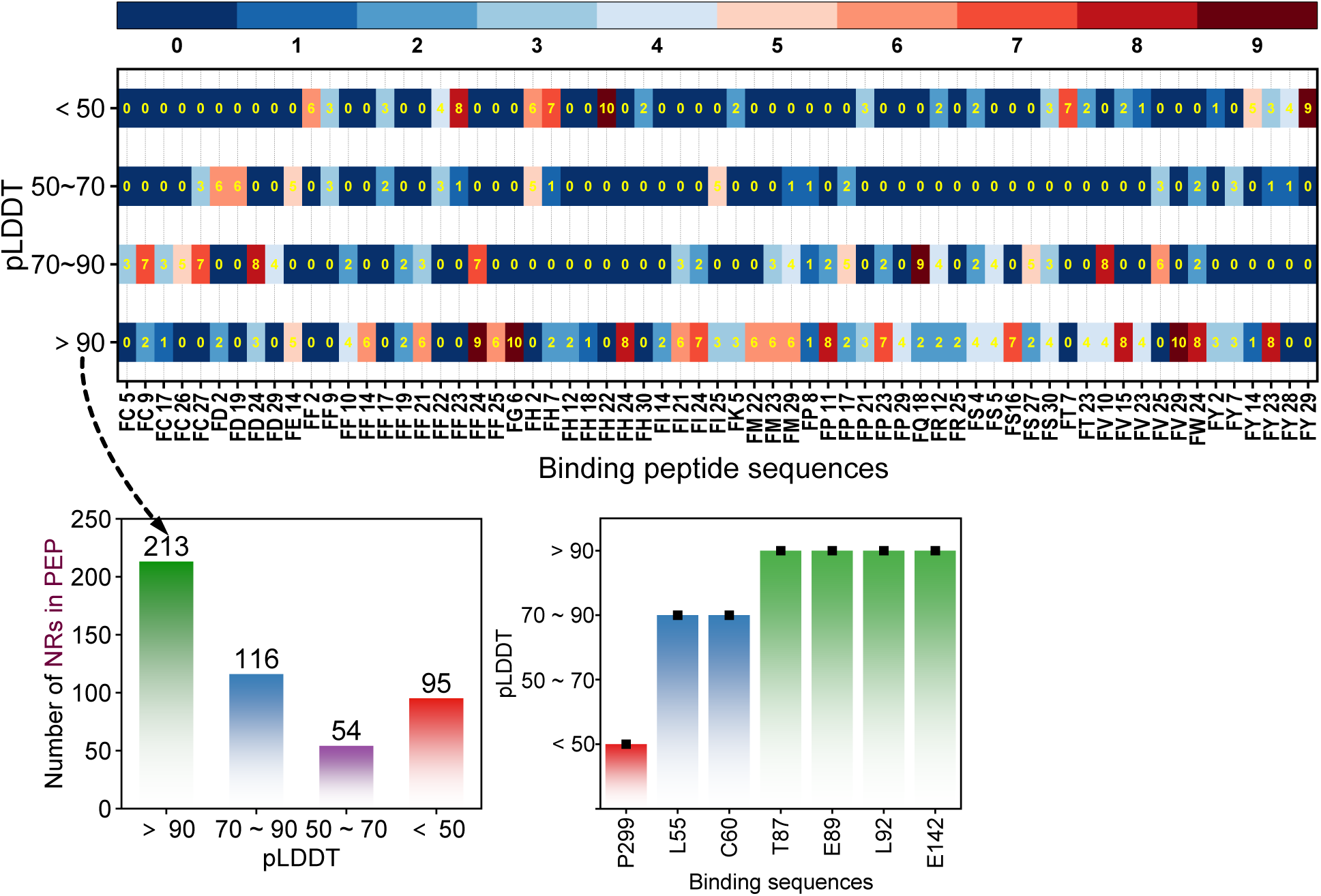
Correlation between pLDDT values and active residues. **(a)** Number of active residues binding to each dipeptides, located within each pLDDT range. **(b)** Total number of active residues within each pLDDT range. The acronym “NRs” in y axis stands for nearby residues. **(c)** pLDDT values of the most active and moderate active residues.

The prevalence of binding activity in low-pLDDT regions (pLDDT < 50) challenges the rigid “lock-and-key” paradigm, ^21, 22^ indicating these likely represent intrinsically disordered regions (IDRs) or flexible loops that adopt structure upon ligand binding via induced fit or conformational selection. This aligns with dynamic binding mechanisms where conformational plasticity enables interactions with diverse ligands. Our data suggest pLDDT may underestimate the functional relevance of such flexible regions due to its emphasis on static structures. Conversely, the concentration of binding residues in high-pLDDT regions (pLDDT > 90) underscores the role of well-defined pockets in molecular recognition, reinforcing pLDDT’s utility for identifying structured sites while highlighting that high confidence alone cannot define functionality, as many such residues are non-interacting.

Collectively, pLDDT operates as a context-dependent indicator: High values (> 70) strongly signal structurally ordered regions where binding typically follows conventional pocket-based mechanisms, while low values (< 50) suggest structural disorder yet do not preclude function, i.e., these regions may mediate transient, dynamic, or allosteric interactions critical for signaling or promiscuous binding. The bimodal distribution in Figure 6b implies pLDDT reliably identifies ordered binding pockets but may overlook functional IDRs. Consequently, relying solely on pLDDT for binding site prediction risks missing flexible interfaces; integrating it with metrics of evolutionary conservation, physicochemical pocket properties, or dynamics-focused algorithms (e.g., MD simulations) could improve functional annotation. The prominence of low-pLDDT binding in FF/FH/FT/FY interactions warrants investigation into whether aromatic dipeptides preferentially target disordered regions through aggregation-induced hydrophobicity mechanism (AIHB). ^23^

## Conclusions

This study establishes the PEP as a viable therapeutic target for cardiovascular diseases by successfully characterizing its dynamic structure and identifying high-affinity dipeptide inhibitors. Initial structure prediction using AF3 revealed intrinsic flexibility, particularly in low-confidence (pLDDT < 50) regions, necessitating refinement via extensive MD simulations. Comparative force field evaluation demonstrated that the AF3-predicted structure experiences significant structural refinement, especially in the low-pLDDT regions. providing an optimized structure for subsequent binding studies. Systematic screening of 20 ‘F-series’ dipeptides (FA-FY) through high-throughput MD simulations identified several stable binders, with FF exhibiting the highest binding propensity (37% of replicates), synergistically driven by hydrogen bonding, hydrophobic interactions, and *π*−*π* stacking. Notably, FF dipeptides concurrently self-assembled into clusters that enhanced PEP binding through multivalent avidity and bridged discontinuous epitopes, revealing an aggregation-dependent mechanism distinct from conventional lock-and-key models. Spatial mapping identified a continuous binding groove involving key residues (e.g., L55, C60, T87, E89, L92, E142), while correlation analysis confirmed functional binding sites exist across both high- and low-pLDDT regions, underscoring the role of conformational plasticity in ligand engagement. These findings validate the therapeutic potential of dipeptides, particularly FF, for selectively inhibiting PEP to mitigate cardiomyocyte apoptosis and ischemia-reperfusion injury, providing a robust computational foundation for future mechanistic studies and peptide-based drug development against cardiovascular pathologies.

## Methods

### Structure Prediction by AF3

The amino acid sequence of the human PHLDA1 protein was retrieved from the UniProt database (website: https://www.uniprot.org/uniprotkb/A2BDE7/entry). The complete protein sequence contains 312 amino acids. This sequence was subsequently submitted to AF3 to generate a predicted three-dimensional protein structure, including the atomic coordinates of all constituent residues.

### Structure Optimization

To assess the structural stability and accuracy of the predicted PHLDA1 model, we conducted all-atom molecular dynamics simulations using GROMACS 2023.2. Simulations were performed in parallel for three times using distinct force fields: AMBER03 for protein with AMBER94 for nucleic acids;^24^ the CHARMM27 all-atom force field;^25^ and AMBER99SB-ILDN.^26^ For each system, the protein structure was solvated within a cubic periodic boundary box (18 × 18 × 18 nm^3^) filled with explicit TIP3P water molecules. Before energy minimization, the systems were charge-neutralized by adding proper amount of Cl*^−^* or Na^+^ counterions. Production simulations comprised sequential equilibration: 20 ns under NVT ensemble conditions (constant number of particle, volume, and temperature = 300 K) followed by 400 ns under NPT ensemble conditions (constant number of particle, temperature = 300 K, pressure = 1 bar). Periodic boundary condition (PBC) was applied in three dimensions. Trajectory after production was processed by applying GROMACS utilities to enforce consistent handling: the protein was centered in the box (-pbc center), translational jumps across PBCs were corrected (-pbc nojump), and overall system rotation/translation was removed via least-squares fitting to the initial structure (-fit rot+trans). Finally, -pbc mol was applied to maintain molecular continuity and ensure the protein remained unconstrained by box boundaries throughout the analyzed trajectory frames.

### Structure Assessment

Given that an accurate structural model is fundamental for structure-based drug design, we assessed the optimized PEP stability by evaluating key kinetic and thermodynamic metrics: root-mean-square deviation (RMSD), radius of gyration (R*_g_*), potential energy, and root-mean-square fluctuation (RMSF). RMSD, Rg, and RMSF, calculated directly from the post-processed trajectories using GROMACS built-in commands. Specifically, the command “gmx rms -s ref.gro -f traj.xtc -o rmsd.xvg” computed RMSD relative to the initial structure. RMSF per residue was generated via “gmx rmsf -s ref.gro -f traj.xtc -o rmsf.xvg”, while R*_g_* was calculated using “gmx gyrate -s topol.tpr -f traj.xtc -o R*_g_*.xvg”. To isolate the protein’s potential energy contribution from the solvated system, a multi-step extraction was performed: First, a protein-only trajectory (prolig.xtc) was created with “gmx trjconv-f full.xtc -s full.tpr -o prolig.xtc”. The corresponding protein topology file (prolig.tpr) was then generated using “gmx convert-tpr -s full.tpr -n prolig.ndx -o prolig.tpr”. This enabled energy recalculation specifically for the protein conformation along the trajectory using “gmx mdrun -s prolig.tpr -rerun prolig.xtc -e prolig.edr”. Finally, the time series of protein potential energy was extracted via “gmx energy -f prolig.edr -o prolig-energy.xvg”. All quantitative plots were generated using OriginPro, while molecular structure figures were rendered in PyMOL.

### MD Simulations of Peptide-Protein Interaction

To investigate dipeptide binding interactions, we performed MD simulations using the CHARMM27 force field-optimized PEP structure, also with CHARMM27 force field. The protein was solvated in a cubic periodic boundary box (10 nm × 10 nm × 10 nm), to which 30 identical dipeptides (representing all 20 standard amino acids: FA, FC, FD, FE, FF, FG, FH, FI, FK, FL, FM, FN, FP, FQ, FR, FS, FT, FV, FW, FY) were added per simulation system. Each system underwent the established protocol: energy minimization followed by 20 ns NVT equilibration and 400 ns NPT production simulation at 300 K and 1 bar. Trajectories were post-processed using identical periodic boundary condition corrections as applied during structural assessment. Peptide-protein complext conformational stability was quantified by calculating RMSD values relative to the initial protein structure (t=0 ns) for all 600 dipeptides (30 peptides × 20 simulations). This involved: 1) Generating peptide-specific index groups (gmx make_ndx -f sys.gro -o peptides.ndx); 2) Extracting the t = 0 ns reference structure of protein-peptide (gmx trjconv -f sys.xtc -s sys.tpr -o ref.gro -b 0 -e 0); 3) Iteratively computing RMSD for each peptide group relative to the protein target using gmx rms -s ref.gro -f sys.xtc -n peptides.ndx -o rmsd-pepX.xvg. The standard deviation of RMSD values during the 200-400 ns trajectory segment was calculated for each dipeptide (totally 600 dipeptides). Peptides exhibiting standard deviation (STD) of RMSD below 0.15 nm were identified as stable binders for subsequent analysis.

## Supporting information

Supplementary RMSD files

## Acknowledgement

Dr. Jiaqi Wang acknowledges the funding support of the National Natural Science Foundation of China (No. 52101023) and Basic Research Program of Jiangsu - General Program of Jiangsu Provincial Department of Science and Technology (No. BK20241816). The authors also acknowledge the high-performance computing platform at Xi’an Jiaotong-Liverpool University and Beijing Paratera Tech Corp. LTD.

