## Supplementary RMSD files for "*In Silico* Screening of Peptide Inhibitors of PHLDA1-Encoded Protein Targeting Cardiovascular Diseases"

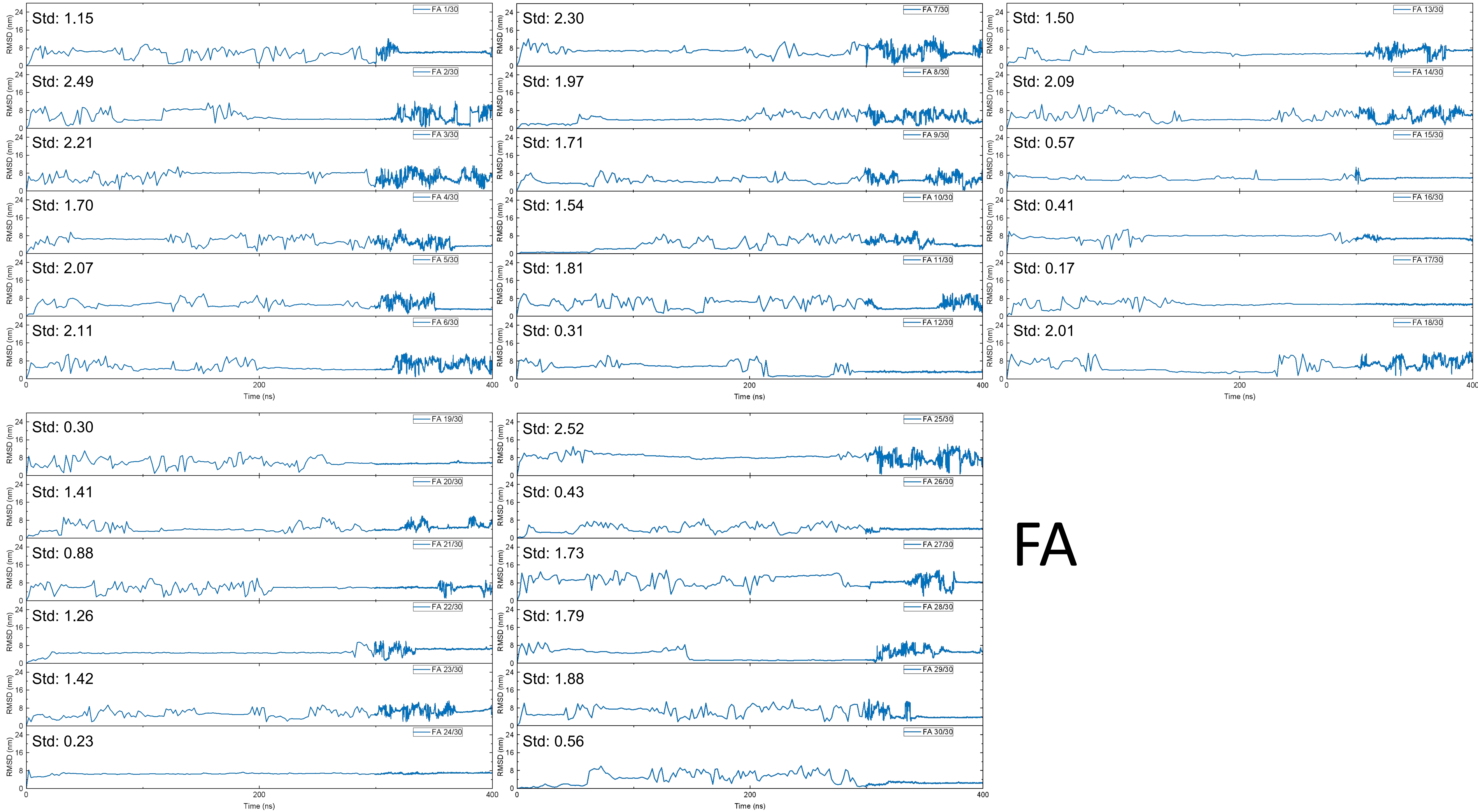
Std: 1.15
FA 1/30
Std: 2.49
FA 2/30
Std: 2.21
FA 3/30
Std: 1.70
FA 4/30
Std: 2.07
FA 5/30
Std: 2.11
FA 6/30
Std: 2.30
FA 7/30
Std: 1.97
FA 8/30
Std: 1.71
FA 9/30
Std: 1.54
FA 10/30
Std: 1.81
FA 11/30
Std: 0.31
FA 12/30
Std: 1.50
FA 13/30
Std: 2.09
FA 14/30
Std: 0.57
FA 15/30
Std: 0.41
FA 16/30
Std: 0.17
FA 17/30
Std: 2.01
FA 18/30
Std: 0.30
FA 19/30
Std: 1.41
FA 20/30
Std: 0.88
FA 21/30
Std: 1.26
FA 22/30
Std: 1.42
FA 23/30
Std: 0.23
FA 24/30
Std: 2.52
FA 25/30
Std: 0.43
FA 26/30
Std: 1.73
FA 27/30
Std: 1.79
FA 28/30
Std: 1.88
FA 29/30
Std: 0.56
FA 30/30
RMSD (nm)
Time (ns)
FA

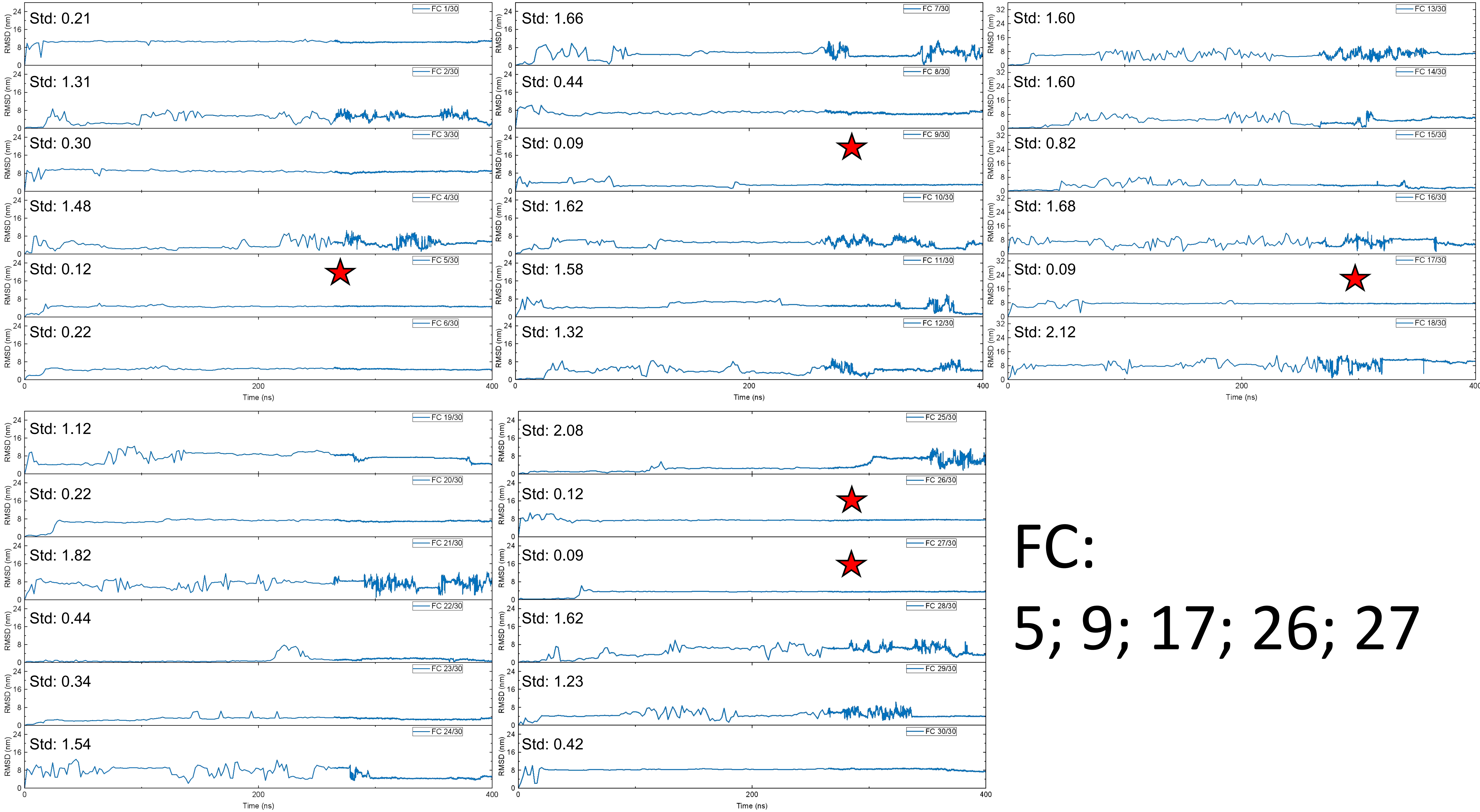

FC 1/30
Std: 0.21
FC 2/30
Std: 1.31
FC 3/30
Std: 0.30
FC 4/30
Std: 1.48
FC 5/30
Std: 0.12
FC 6/30
Std: 0.22
FC 7/30
Std: 1.66
FC 8/30
Std: 0.44
FC 9/30
Std: 0.09
FC 10/30
Std: 1.62
FC 11/30
Std: 1.58
FC 12/30
Std: 1.32
FC 13/30
Std: 1.60
FC 14/30
Std: 1.60
FC 15/30
Std: 0.82
FC 16/30
Std: 1.68
FC 17/30
Std: 0.09
FC 18/30
Std: 2.12
FC 19/30
Std: 1.12
FC 20/30
Std: 0.22
FC 21/30
Std: 1.82
FC 22/30
Std: 0.44
FC 23/30
Std: 0.34
FC 24/30
Std: 1.54
FC 25/30
Std: 2.08
FC 26/30
Std: 0.12
FC 27/30
Std: 0.09
FC 28/30
Std: 1.62
FC 29/30
Std: 1.23
FC 30/30
Std: 0.42
RMSD (nm)
Time (ns)


FC:

5; 9; 17; 26; 27

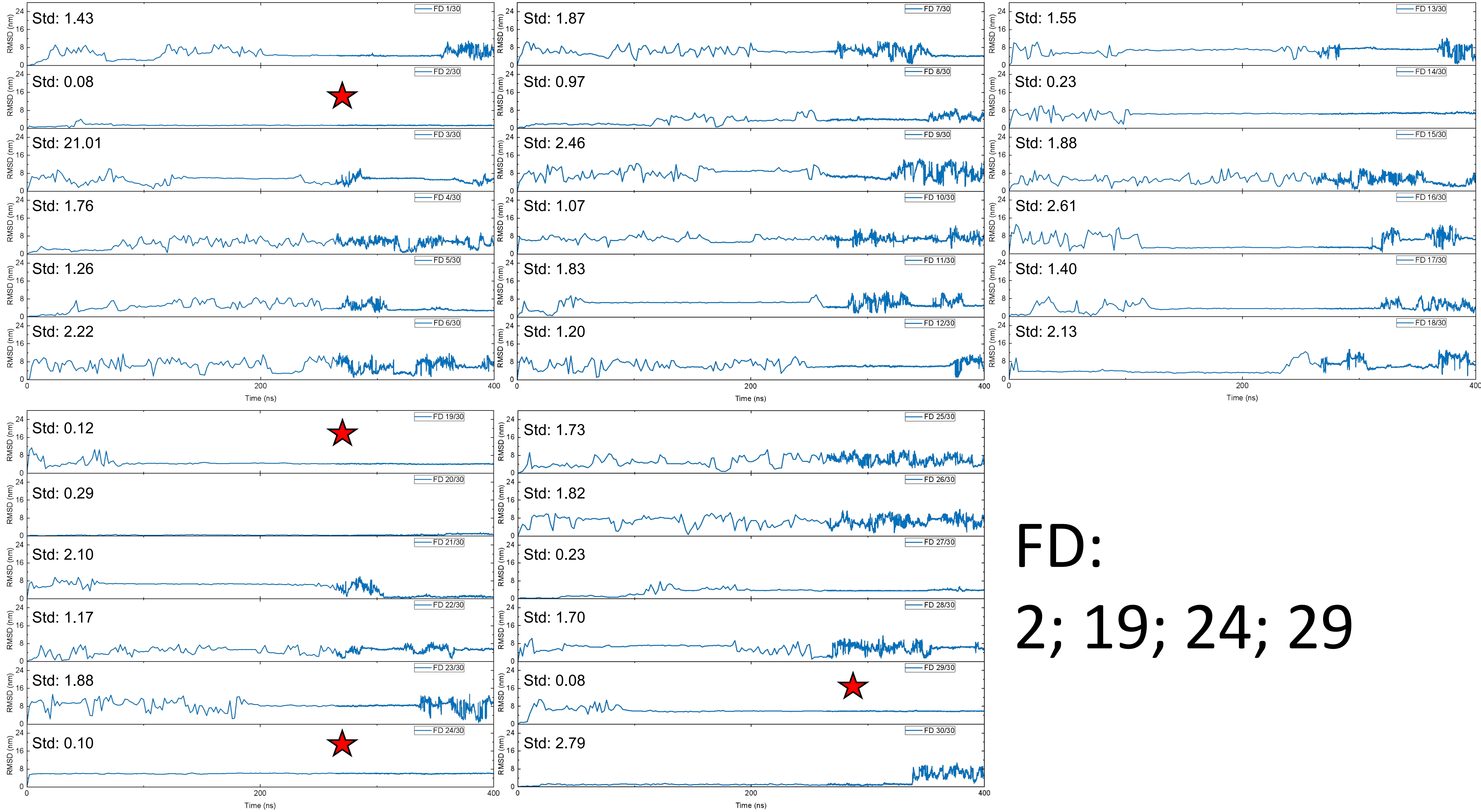

FD 1/30
Std: 1.43
FD 2/30
Std: 0.08
FD 3/30
Std: 21.01
FD 4/30
Std: 1.76
FD 5/30
Std: 1.26
FD 6/30
Std: 2.22
FD 7/30
Std: 1.87
FD 8/30
Std: 0.97
FD 9/30
Std: 2.46
FD 10/30
Std: 1.07
FD 11/30
Std: 1.83
FD 12/30
Std: 1.20
FD 13/30
Std: 1.55
FD 14/30
Std: 0.23
FD 15/30
Std: 1.88
FD 16/30
Std: 2.61
FD 17/30
Std: 1.40
FD 18/30
Std: 2.13
FD 19/30
Std: 0.12
FD 20/30
Std: 0.29
FD 21/30
Std: 2.10
FD 22/30
Std: 1.17
FD 23/30
Std: 1.88
FD 24/30
Std: 0.10
FD 25/30
Std: 1.73
FD 26/30
Std: 1.82
FD 27/30
Std: 0.23
FD 28/30
Std: 1.70
FD 29/30
Std: 0.08
FD 30/30
Std: 2.79
RMSD (nm)
Time (ns)
FD:
2; 19; 24; 29

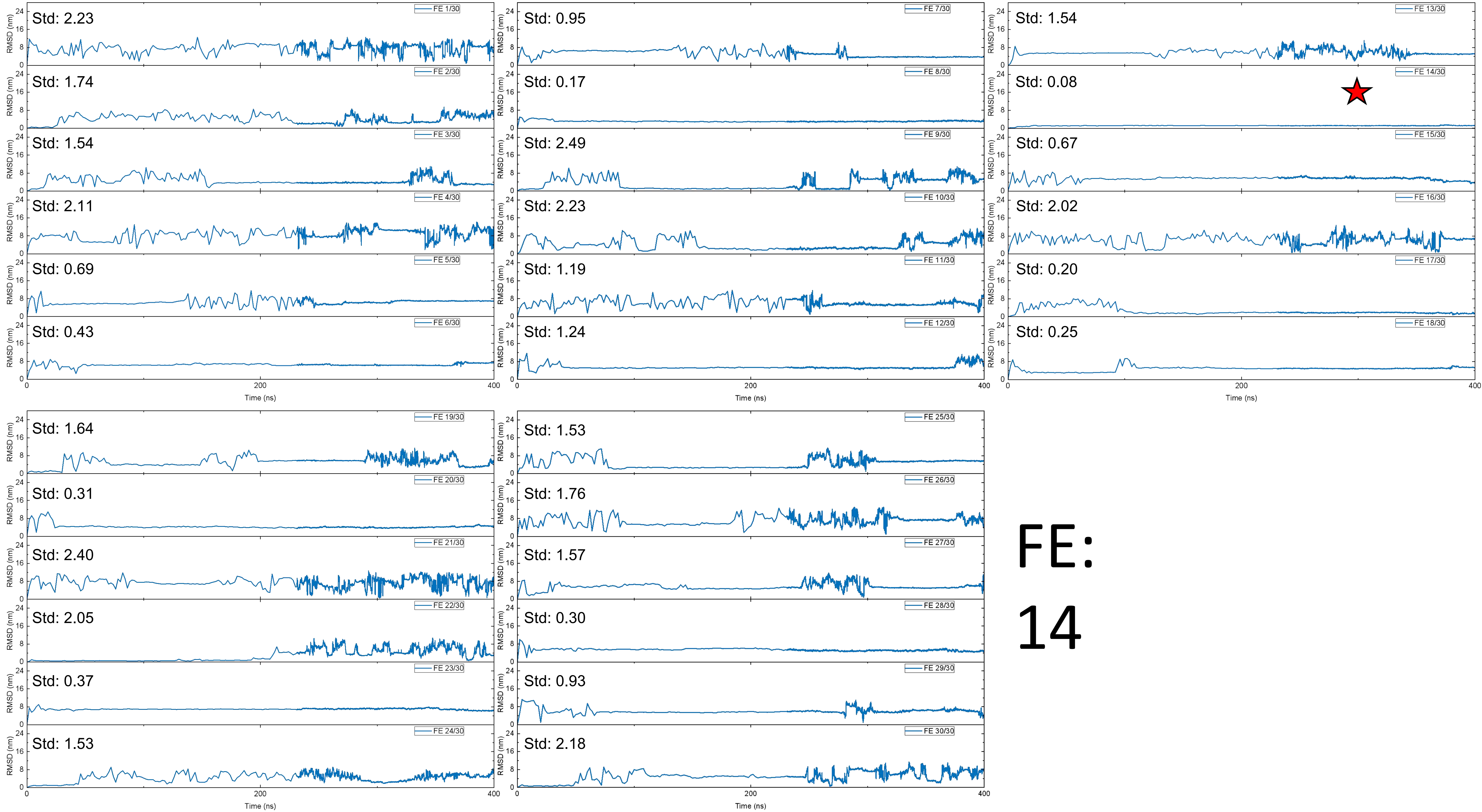
FE 1/30
Std: 2.23
FE 2/30
Std: 1.74
FE 3/30
Std: 1.54
FE 4/30
Std: 2.11
FE 5/30
Std: 0.69
FE 6/30
Std: 0.43
FE 7/30
Std: 0.95
FE 8/30
Std: 0.17
FE 9/30
Std: 2.49
FE 10/30
Std: 2.23
FE 11/30
Std: 1.19
FE 12/30
Std: 1.24
FE 13/30
Std: 1.54
FE 14/30
Std: 0.08
FE 15/30
Std: 0.67
FE 16/30
Std: 2.02
FE 17/30
Std: 0.20
FE 18/30
Std: 0.25
FE 19/30
Std: 1.64
FE 20/30
Std: 0.31
FE 21/30
Std: 2.40
FE 22/30
Std: 2.05
FE 23/30
Std: 0.37
FE 24/30
Std: 1.53
FE 25/30
Std: 1.53
FE 26/30
Std: 1.76
FE 27/30
Std: 1.57
FE 28/30
Std: 0.30
FE 29/30
Std: 0.93
FE 30/30
Std: 2.18
RMSD (nm)
Time (ns)
FE:
14

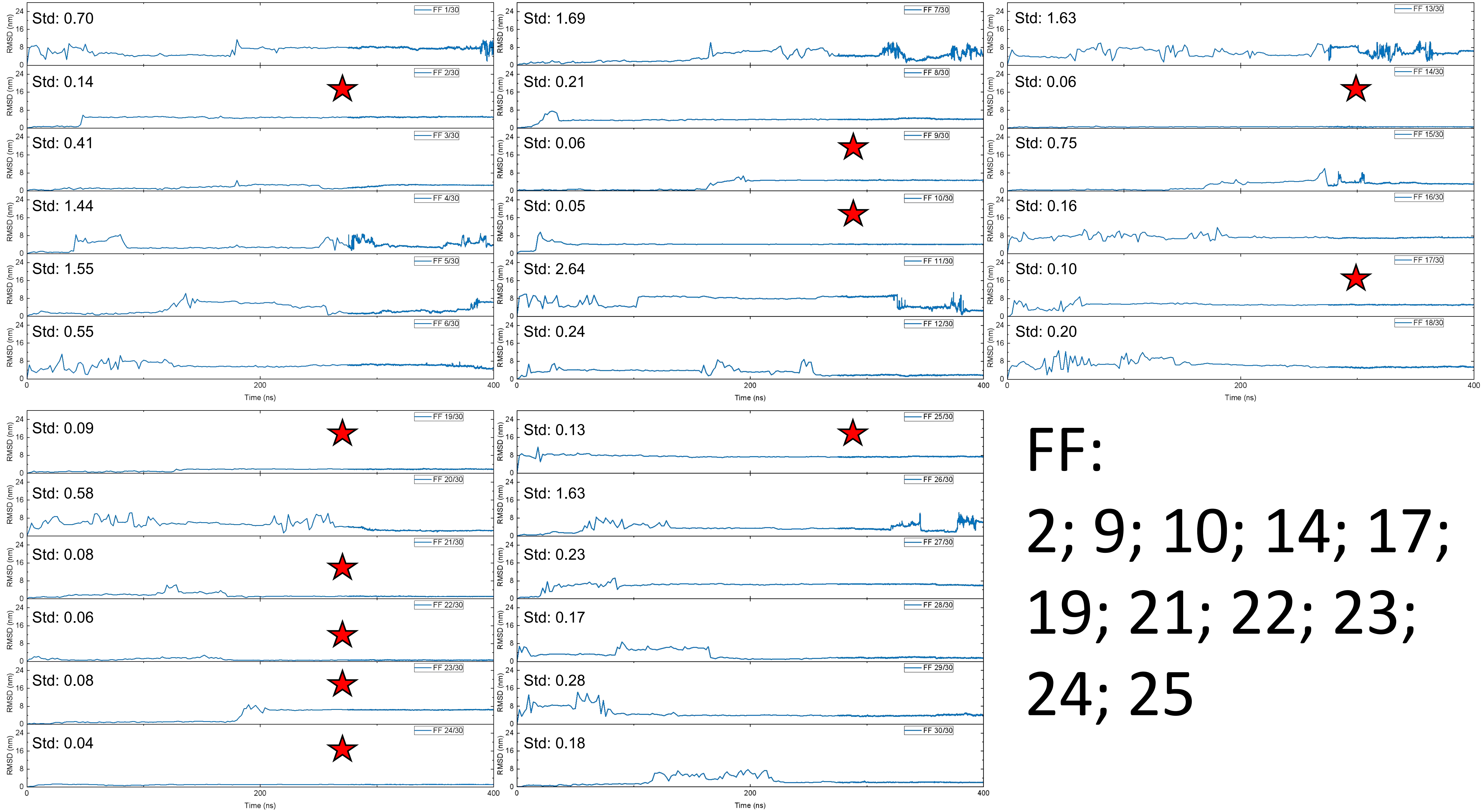
FF 1/30
Std: 0.70
FF 2/30
Std: 0.14
FF 3/30
Std: 0.41
FF 4/30
Std: 1.44
FF 5/30
Std: 1.55
FF 6/30
Std: 0.55
FF 7/30
Std: 1.69
FF 8/30
Std: 0.21
FF 9/30
Std: 0.06
FF 10/30
Std: 0.05
FF 11/30
Std: 2.64
FF 12/30
Std: 0.24
FF 13/30
Std: 1.63
FF 14/30
Std: 0.06
FF 15/30
Std: 0.75
FF 16/30
Std: 0.16
FF 17/30
Std: 0.10
FF 18/30
Std: 0.20
FF 19/30
Std: 0.09
FF 20/30
Std: 0.58
FF 21/30
Std: 0.08
FF 22/30
Std: 0.06
FF 23/30
Std: 0.08
FF 24/30
Std: 0.04
FF 25/30
Std: 0.13
FF 26/30
Std: 1.63
FF 27/30
Std: 0.23
FF 28/30
Std: 0.17
FF 29/30
Std: 0.28
FF 30/30
Std: 0.18
RMSD (nm)
Time (ns)
FF:
2; 9; 10; 14; 17;
19; 21; 22; 23;
24; 25

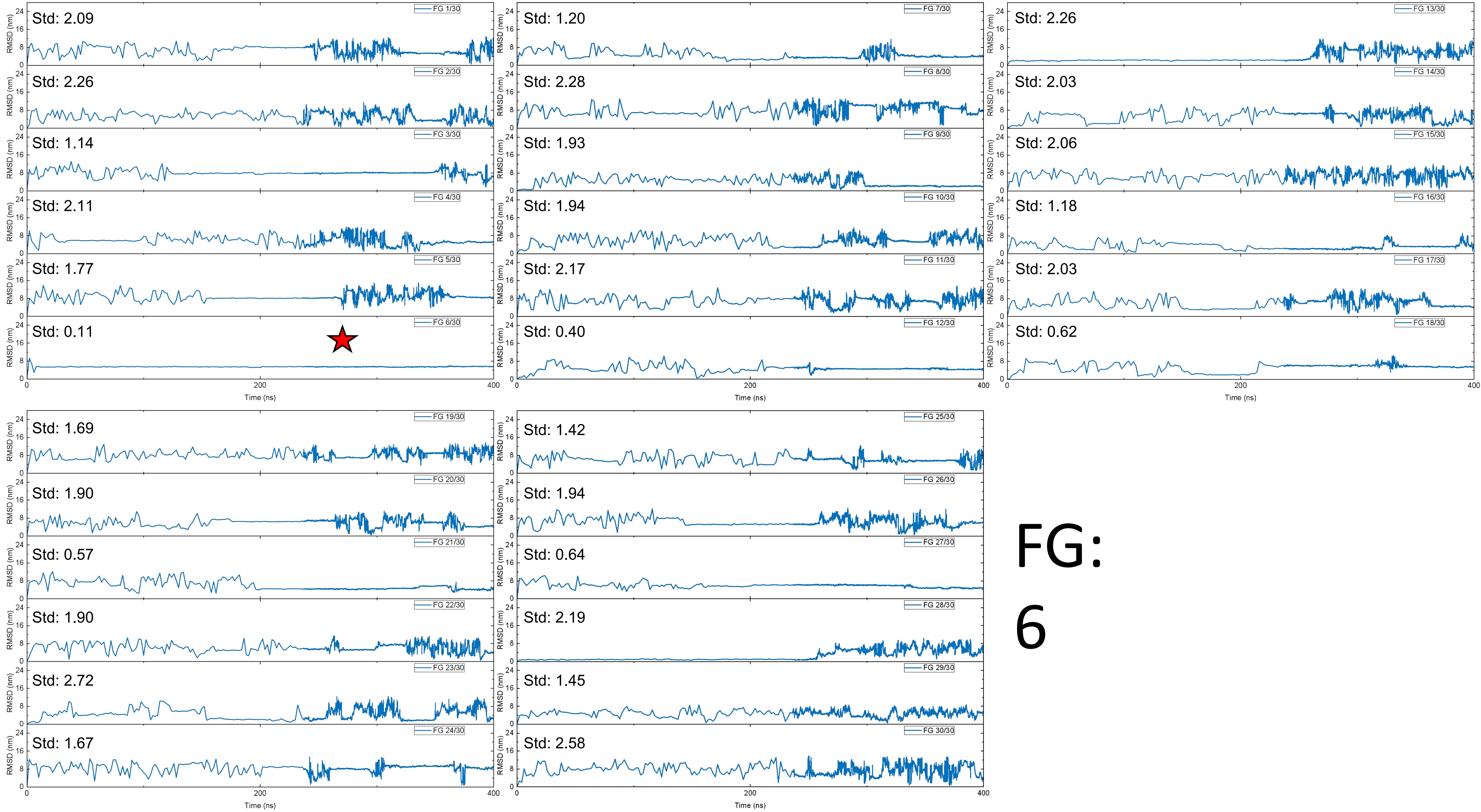
FG 1/30
Std: 2.09
FG 2/30
Std: 2.26
FG 3/30
Std: 1.14
FG 4/30
Std: 2.11
FG 5/30
Std: 1.77
FG 6/30
Std: 0.11
FG 7/30
Std: 1.20
FG 8/30
Std: 2.28
FG 9/30
Std: 1.93
FG 10/30
Std: 1.94
FG 11/30
Std: 2.17
FG 12/30
Std: 0.40
FG 13/30
Std: 2.26
FG 14/30
Std: 2.03
FG 15/30
Std: 2.06
FG 16/30
Std: 1.18
FG 17/30
Std: 2.03
FG 18/30
Std: 0.62
FG 19/30
Std: 1.69
FG 20/30
Std: 1.90
FG 21/30
Std: 0.57
FG 22/30
Std: 1.90
FG 23/30
Std: 2.72
FG 24/30
Std: 1.67
FG 25/30
Std: 1.42
FG 26/30
Std: 1.94
FG 27/30
Std: 0.64
FG 28/30
Std: 2.19
FG 29/30
Std: 1.45
FG 30/30
Std: 2.58
RMSD (nm)
Time (ns)
0
8
16
24
200
400

FG:
6

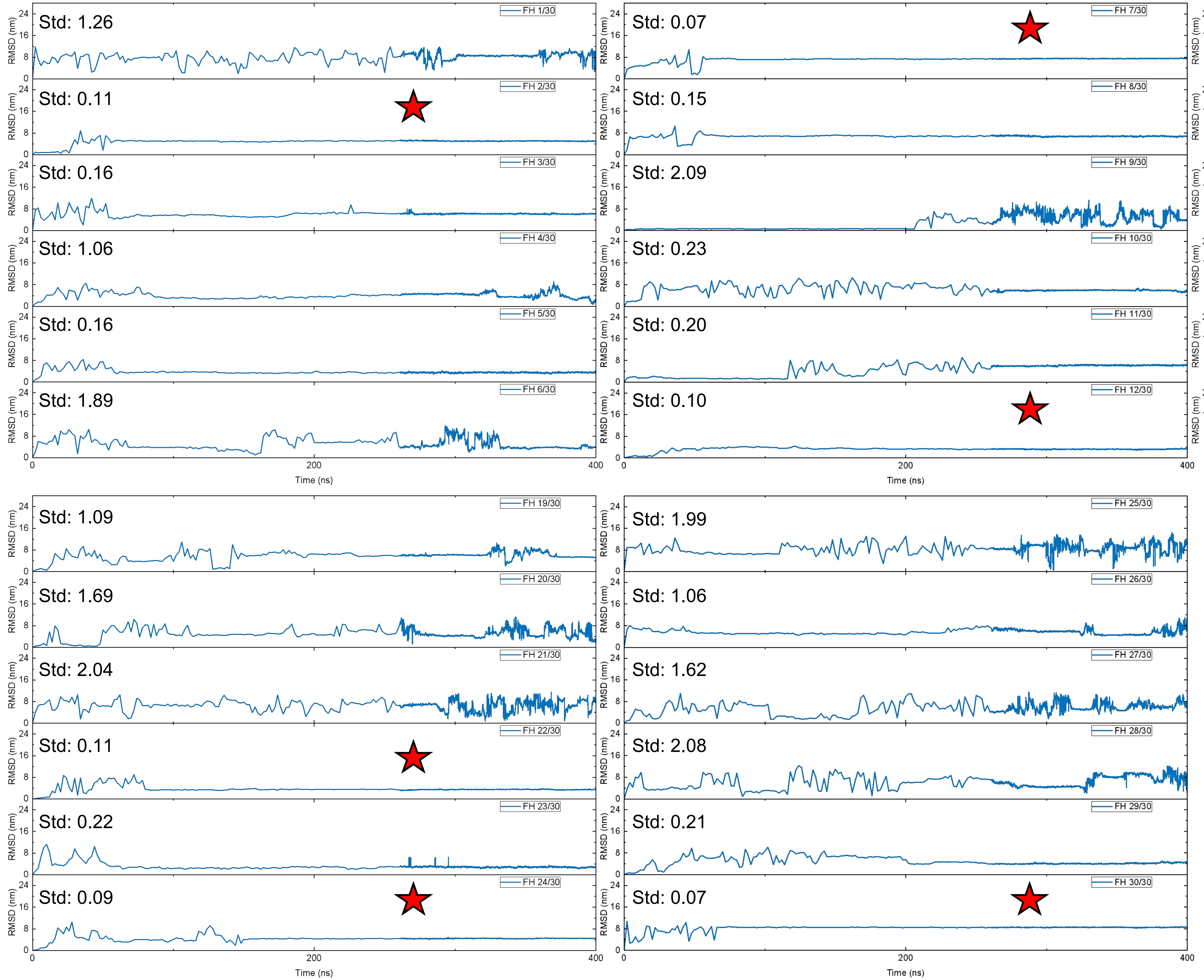


# FH:

# 2; 7; 12; 18; 22; 24; 30

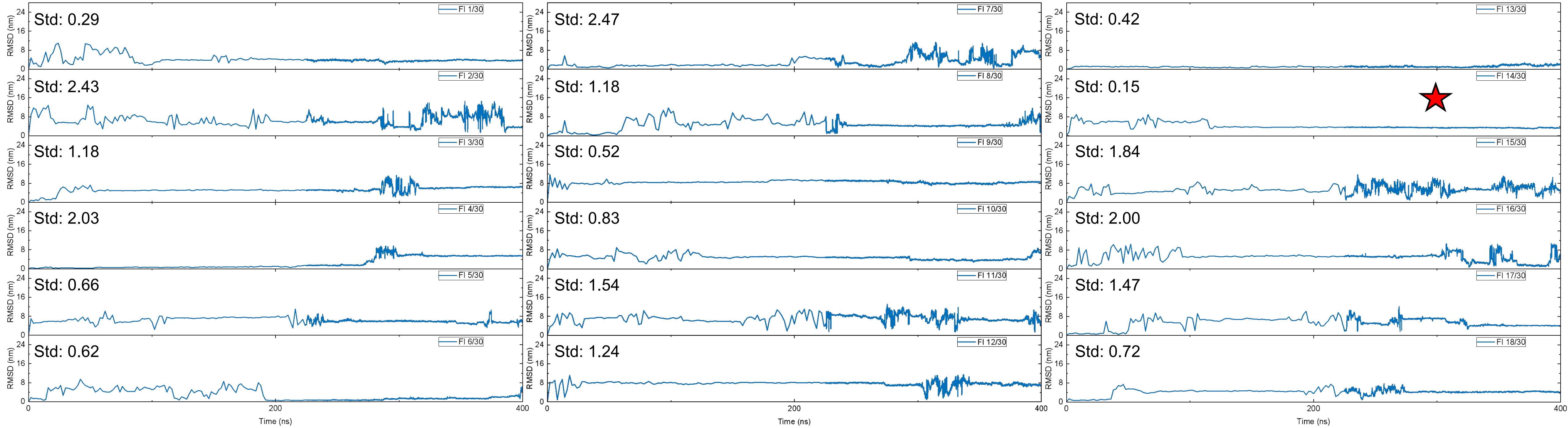
FI 1/30 Std: 0.29
FI 2/30 Std: 2.43
FI 3/30 Std: 1.18
FI 4/30 Std: 2.03
FI 5/30 Std: 0.66
FI 6/30 Std: 0.62
FI 7/30 Std: 2.47
FI 8/30 Std: 1.18
FI 9/30 Std: 0.52
FI 10/30 Std: 0.83
FI 11/30 Std: 1.54
FI 12/30 Std: 1.24
FI 13/30 Std: 0.42
FI 14/30 Std: 0.15
FI 15/30 Std: 1.84
FI 16/30 Std: 2.00
FI 17/30 Std: 1.47
FI 18/30 Std: 0.72
RMSD (nm)
Time (ns)
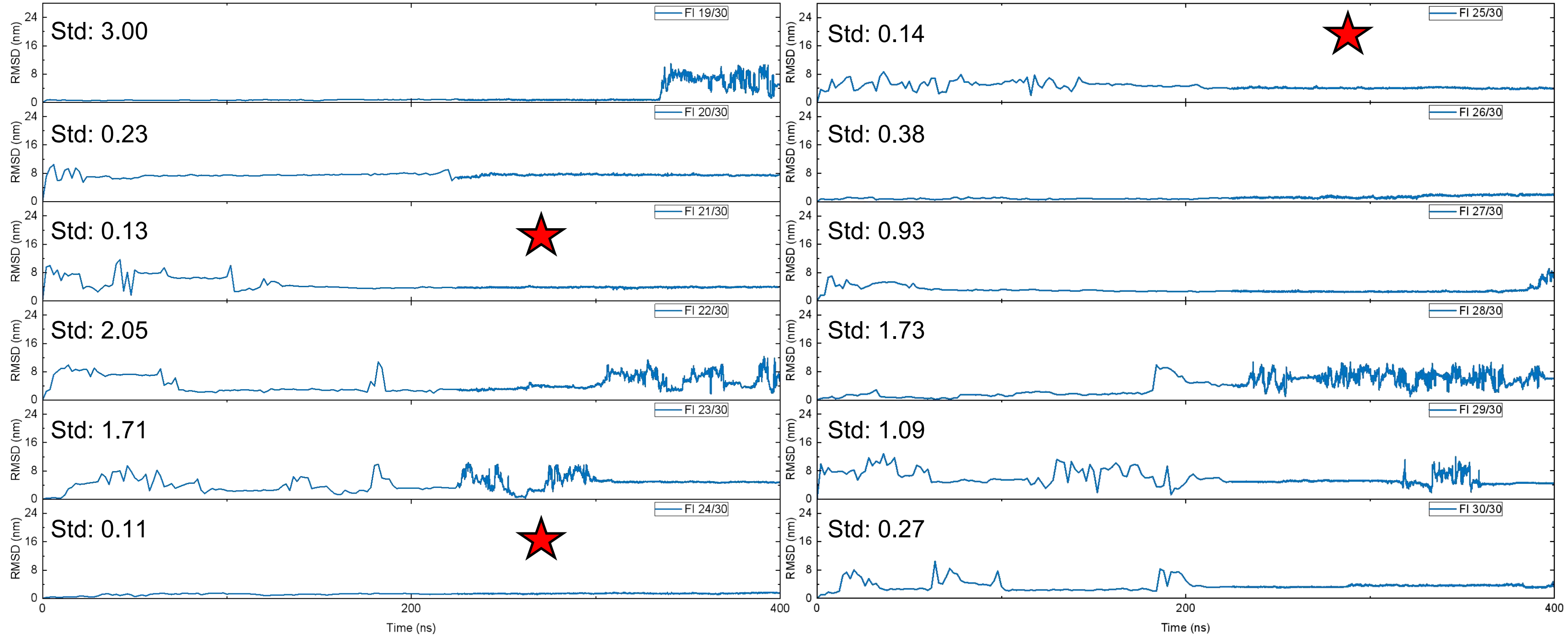
FI 19/30 Std: 3.00
FI 20/30 Std: 0.23
FI 21/30 Std: 0.13
FI 22/30 Std: 2.05
FI 23/30 Std: 1.71
FI 24/30 Std: 0.11
FI 25/30 Std: 0.14
FI 26/30 Std: 0.38
FI 27/30 Std: 0.93
FI 28/30 Std: 1.73
FI 29/30 Std: 1.09
FI 30/30 Std: 0.27
RMSD (nm)
Time (ns)

# FI:

# 14; 21; 24; 25

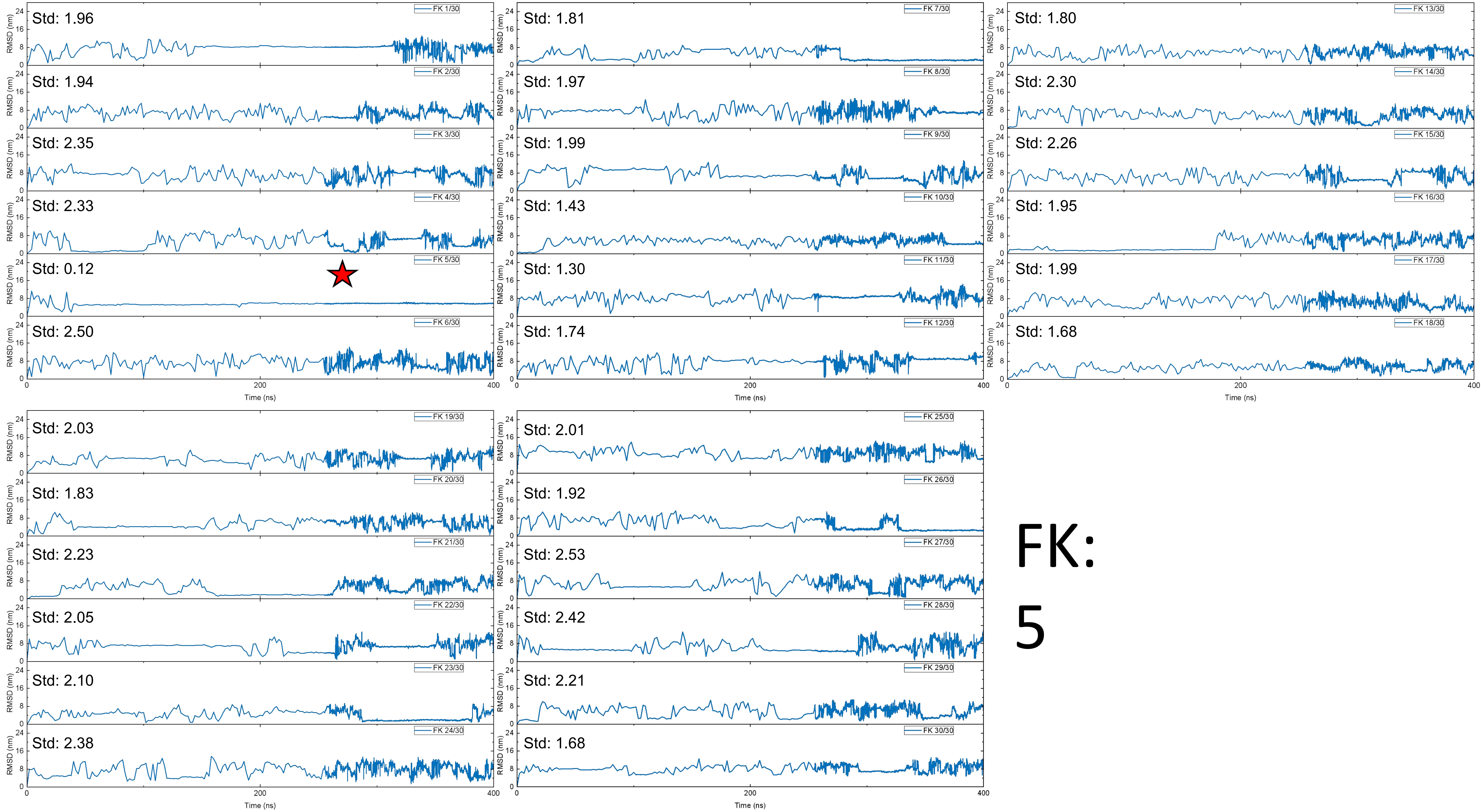
Std: 1.96
FK 1/30
Std: 1.94
FK 2/30
Std: 2.35
FK 3/30
Std: 2.33
FK 4/30
Std: 0.12
FK 5/30
Std: 2.50
FK 6/30
Std: 1.81
FK 7/30
Std: 1.97
FK 8/30
Std: 1.99
FK 9/30
Std: 1.43
FK 10/30
Std: 1.30
FK 11/30
Std: 1.74
FK 12/30
Std: 1.80
FK 13/30
Std: 2.30
FK 14/30
Std: 2.26
FK 15/30
Std: 1.95
FK 16/30
Std: 1.99
FK 17/30
Std: 1.68
FK 18/30
Std: 2.03
FK 19/30
Std: 1.83
FK 20/30
Std: 2.23
FK 21/30
Std: 2.05
FK 22/30
Std: 2.10
FK 23/30
Std: 2.38
FK 24/30
Std: 2.01
FK 25/30
Std: 1.92
FK 26/30
Std: 2.53
FK 27/30
Std: 2.42
FK 28/30
Std: 2.21
FK 29/30
Std: 1.68
FK 30/30
RMSD (nm)
Time (ns)
FK:
5

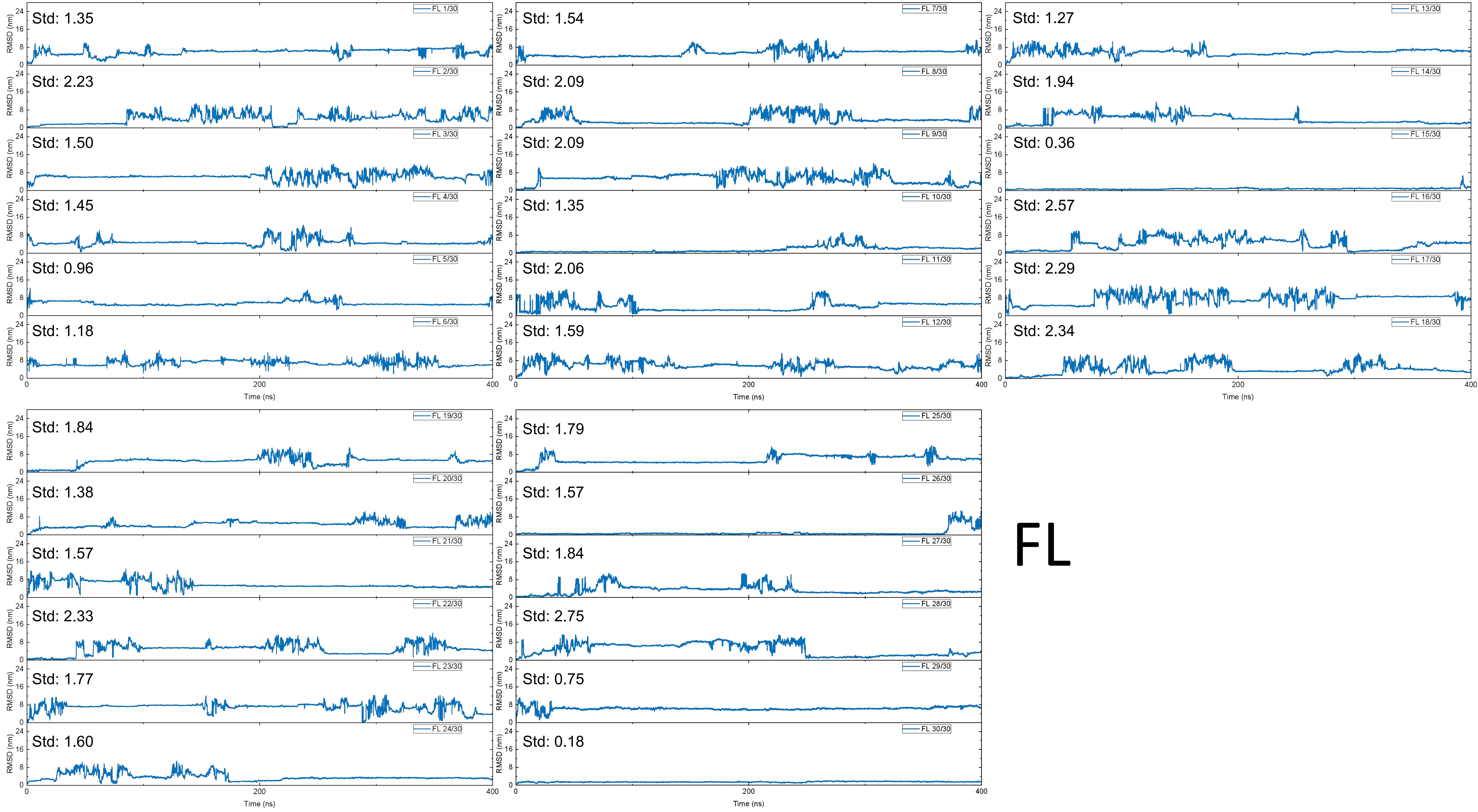
FL
FL 1/30
Std: 1.35
FL 2/30
Std: 2.23
FL 3/30
Std: 1.50
FL 4/30
Std: 1.45
FL 5/30
Std: 0.96
FL 6/30
Std: 1.18
FL 7/30
Std: 1.54
FL 8/30
Std: 2.09
FL 9/30
Std: 2.09
FL 10/30
Std: 1.35
FL 11/30
Std: 2.06
FL 12/30
Std: 1.59
FL 13/30
Std: 1.27
FL 14/30
Std: 1.94
FL 15/30
Std: 0.36
FL 16/30
Std: 2.57
FL 17/30
Std: 2.29
FL 18/30
Std: 2.34
FL 19/30
Std: 1.84
FL 20/30
Std: 1.38
FL 21/30
Std: 1.57
FL 22/30
Std: 2.33
FL 23/30
Std: 1.77
FL 24/30
Std: 1.60
FL 25/30
Std: 1.79
FL 26/30
Std: 1.57
FL 27/30
Std: 1.84
FL 28/30
Std: 2.75
FL 29/30
Std: 0.75
FL 30/30
Std: 0.18
RMSD (nm)
Time (ns)

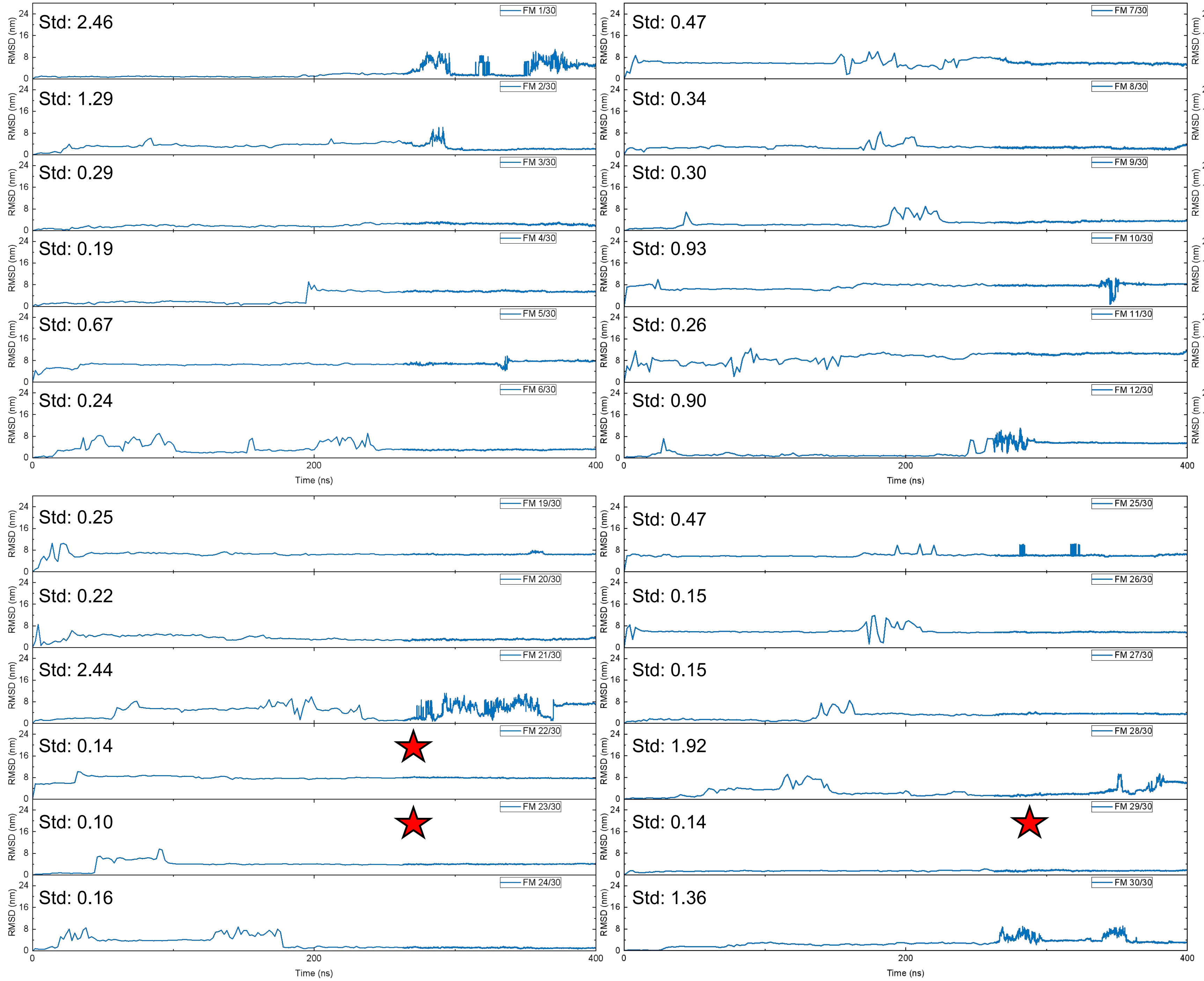
FM 1/30
Std: 2.46
FM 2/30
Std: 1.29
FM 3/30
Std: 0.29
FM 4/30
Std: 0.19
FM 5/30
Std: 0.67
FM 6/30
Std: 0.24
FM 7/30
Std: 0.47
FM 8/30
Std: 0.34
FM 9/30
Std: 0.30
FM 10/30
Std: 0.93
FM 11/30
Std: 0.26
FM 12/30
Std: 0.90
FM 13/30
Std: 0.24
FM 14/30
Std: 0.47
FM 15/30
Std: 0.40
FM 16/30
Std: 0.20
FM 17/30
Std: 0.55
FM 18/30
Std: 0.28
FM 19/30
Std: 0.25
FM 20/30
Std: 0.22
FM 21/30
Std: 2.44
FM 22/30
Std: 0.14
FM 23/30
Std: 0.10
FM 24/30
Std: 0.16
FM 25/30
Std: 0.47
FM 26/30
Std: 0.15
FM 27/30
Std: 0.15
FM 28/30
Std: 1.92
FM 29/30
Std: 0.14
FM 30/30
Std: 1.36
RMSD (nm)
Time (ns)

FM:

22; 23; 29

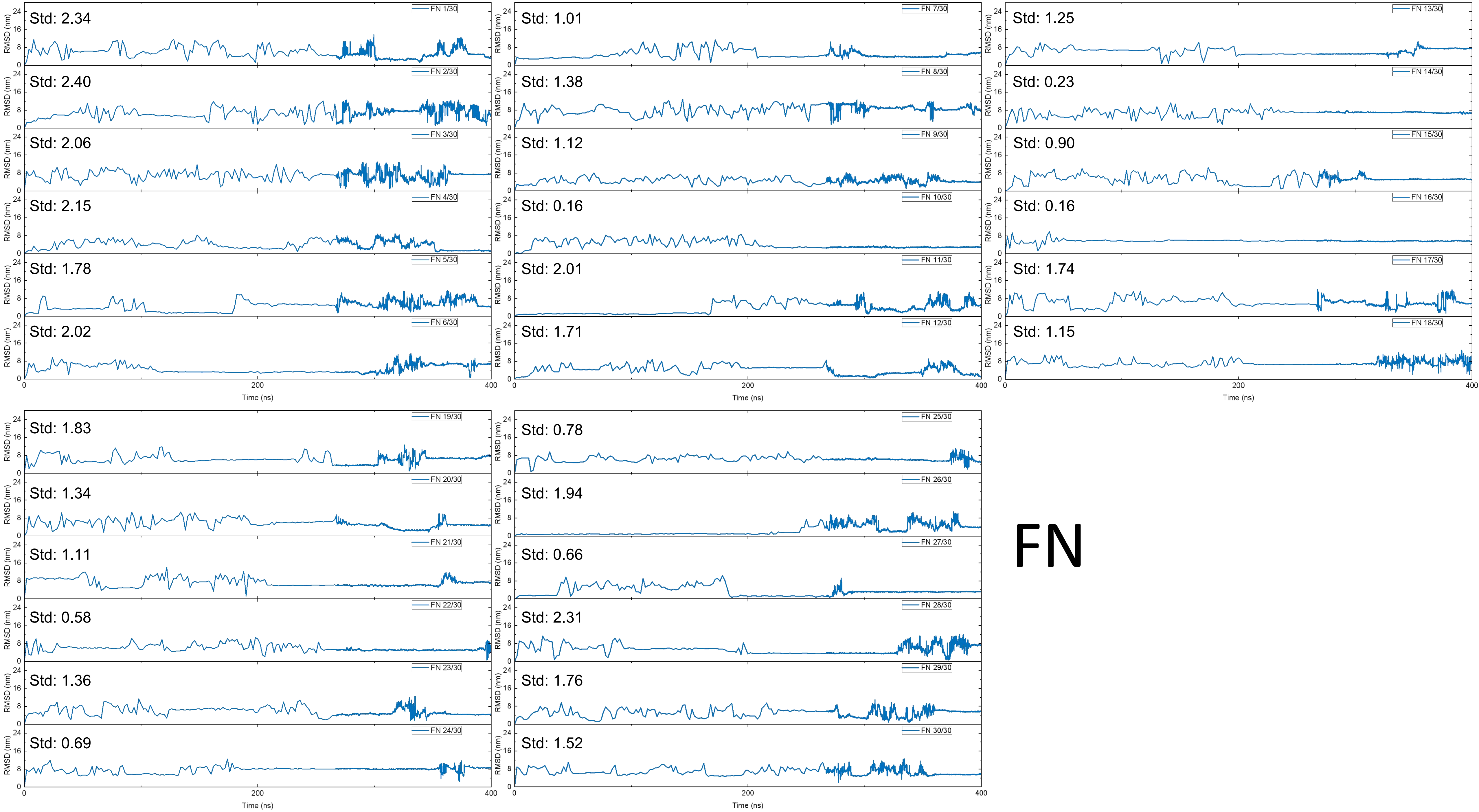

Std: 2.34
FN 1/30
Std: 2.40
FN 2/30
Std: 2.06
FN 3/30
Std: 2.15
FN 4/30
Std: 1.78
FN 5/30
Std: 2.02
FN 6/30
Std: 1.01
FN 7/30
Std: 1.38
FN 8/30
Std: 1.12
FN 9/30
Std: 0.16
FN 10/30
Std: 2.01
FN 11/30
Std: 1.71
FN 12/30
Std: 1.25
FN 13/30
Std: 0.23
FN 14/30
Std: 0.90
FN 15/30
Std: 0.16
FN 16/30
Std: 1.74
FN 17/30
Std: 1.15
FN 18/30
Std: 1.83
FN 19/30
Std: 1.34
FN 20/30
Std: 1.11
FN 21/30
Std: 0.58
FN 22/30
Std: 1.36
FN 23/30
Std: 0.69
FN 24/30
Std: 0.78
FN 25/30
Std: 1.94
FN 26/30
Std: 0.66
FN 27/30
Std: 2.31
FN 28/30
Std: 1.76
FN 29/30
Std: 1.52
FN 30/30
RMSD (nm)
Time (ns)
FN

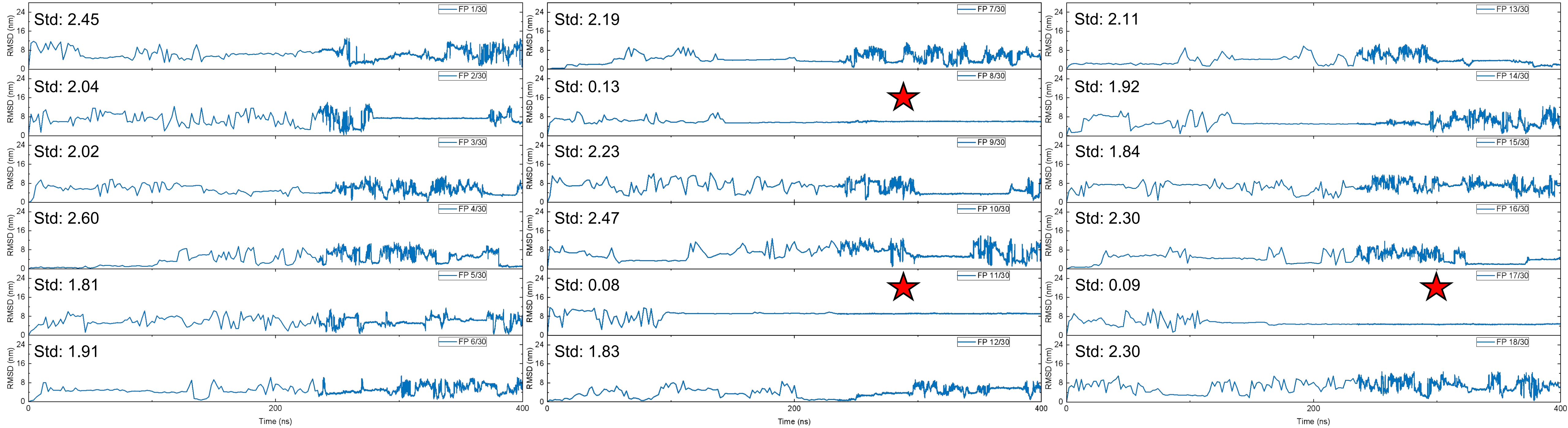


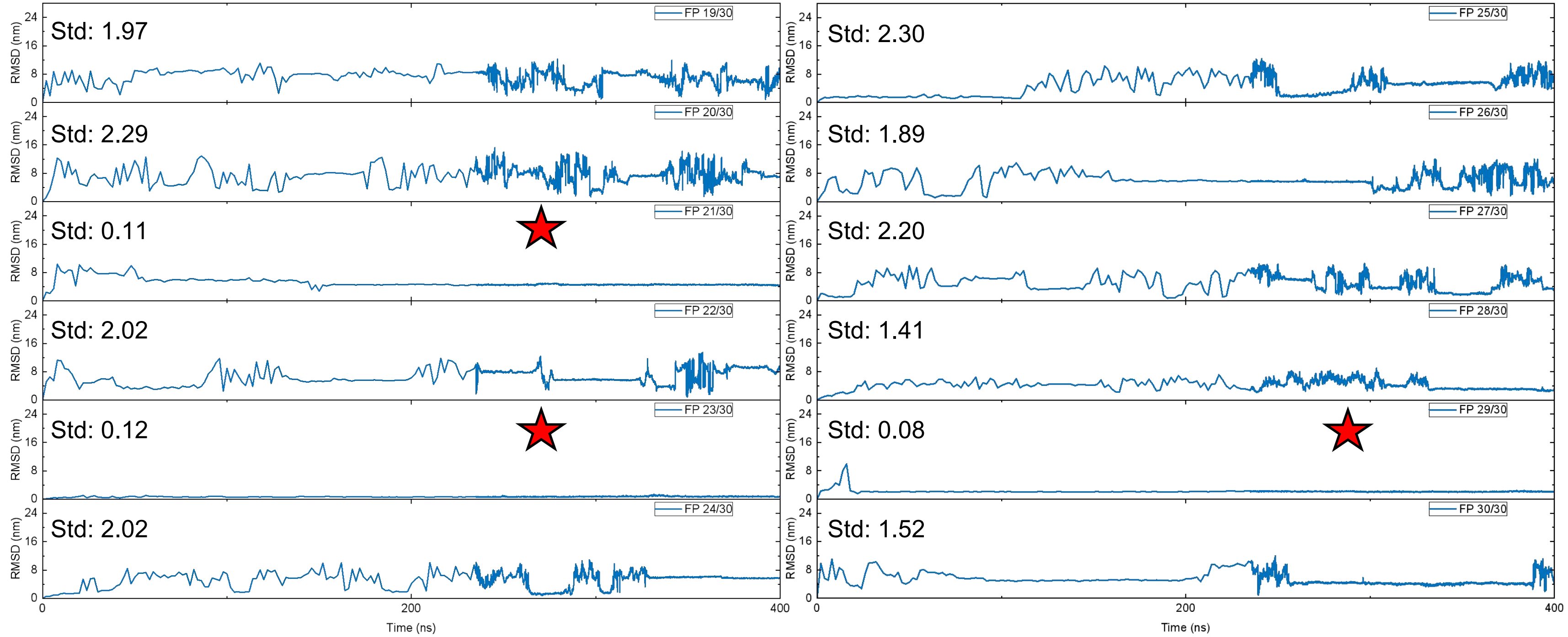


FP:

8; 11; 17; 21; 23; 29

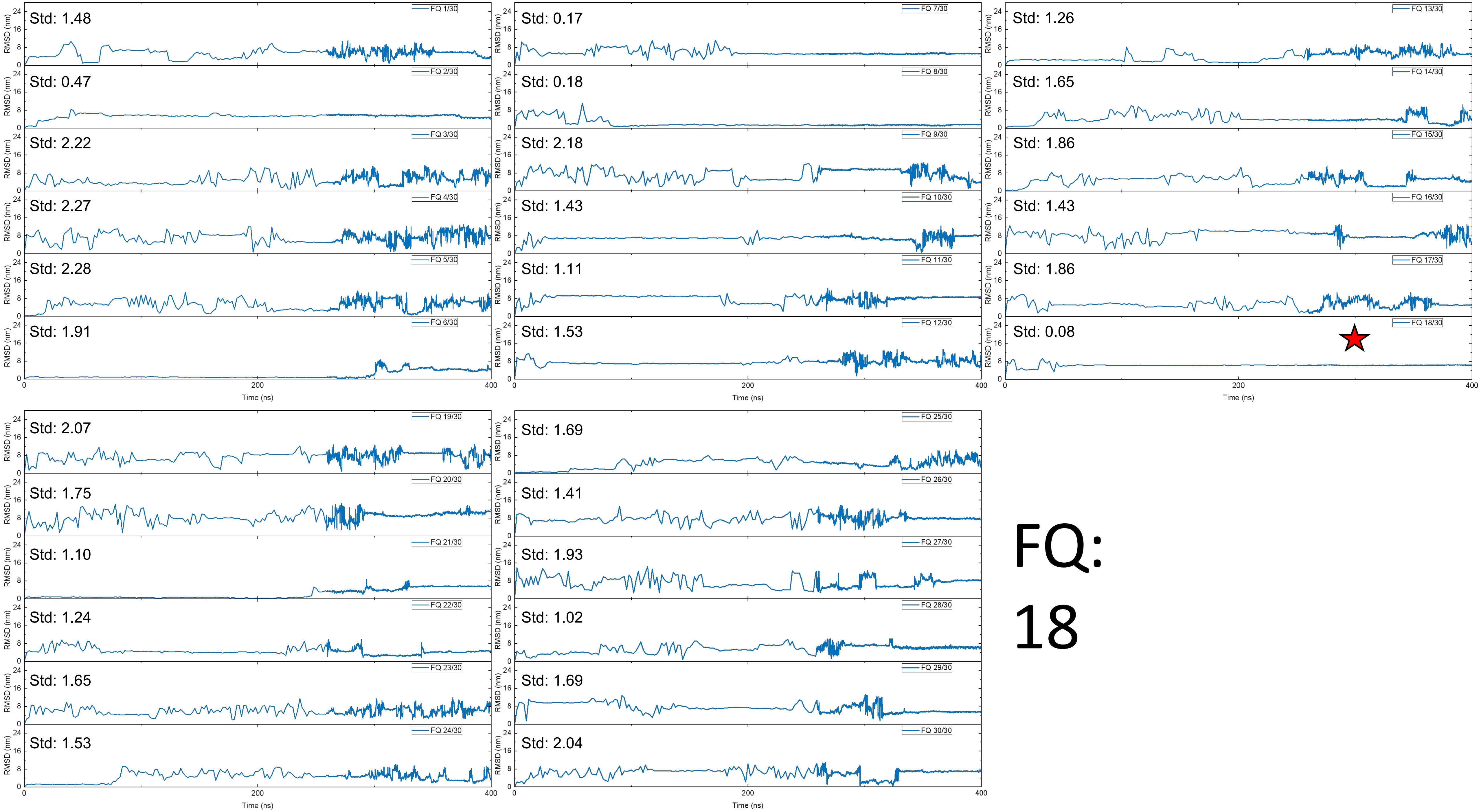

Std: 1.48
FQ 1/30
Std: 0.47
FQ 2/30
Std: 2.22
FQ 3/30
Std: 2.27
FQ 4/30
Std: 2.28
FQ 5/30
Std: 1.91
FQ 6/30
Std: 0.17
FQ 7/30
Std: 0.18
FQ 8/30
Std: 2.18
FQ 9/30
Std: 1.43
FQ 10/30
Std: 1.11
FQ 11/30
Std: 1.53
FQ 12/30
Std: 1.26
FQ 13/30
Std: 1.65
FQ 14/30
Std: 1.86
FQ 15/30
Std: 1.43
FQ 16/30
Std: 1.86
FQ 17/30
Std: 0.08
FQ 18/30
Std: 2.07
FQ 19/30
Std: 1.75
FQ 20/30
Std: 1.10
FQ 21/30
Std: 1.24
FQ 22/30
Std: 1.65
FQ 23/30
Std: 1.53
FQ 24/30
Std: 1.69
FQ 25/30
Std: 1.41
FQ 26/30
Std: 1.93
FQ 27/30
Std: 1.02
FQ 28/30
Std: 1.69
FQ 29/30
Std: 2.04
FQ 30/30
RMSD (nm)
Time (ns)
0
200
400
FQ:
18

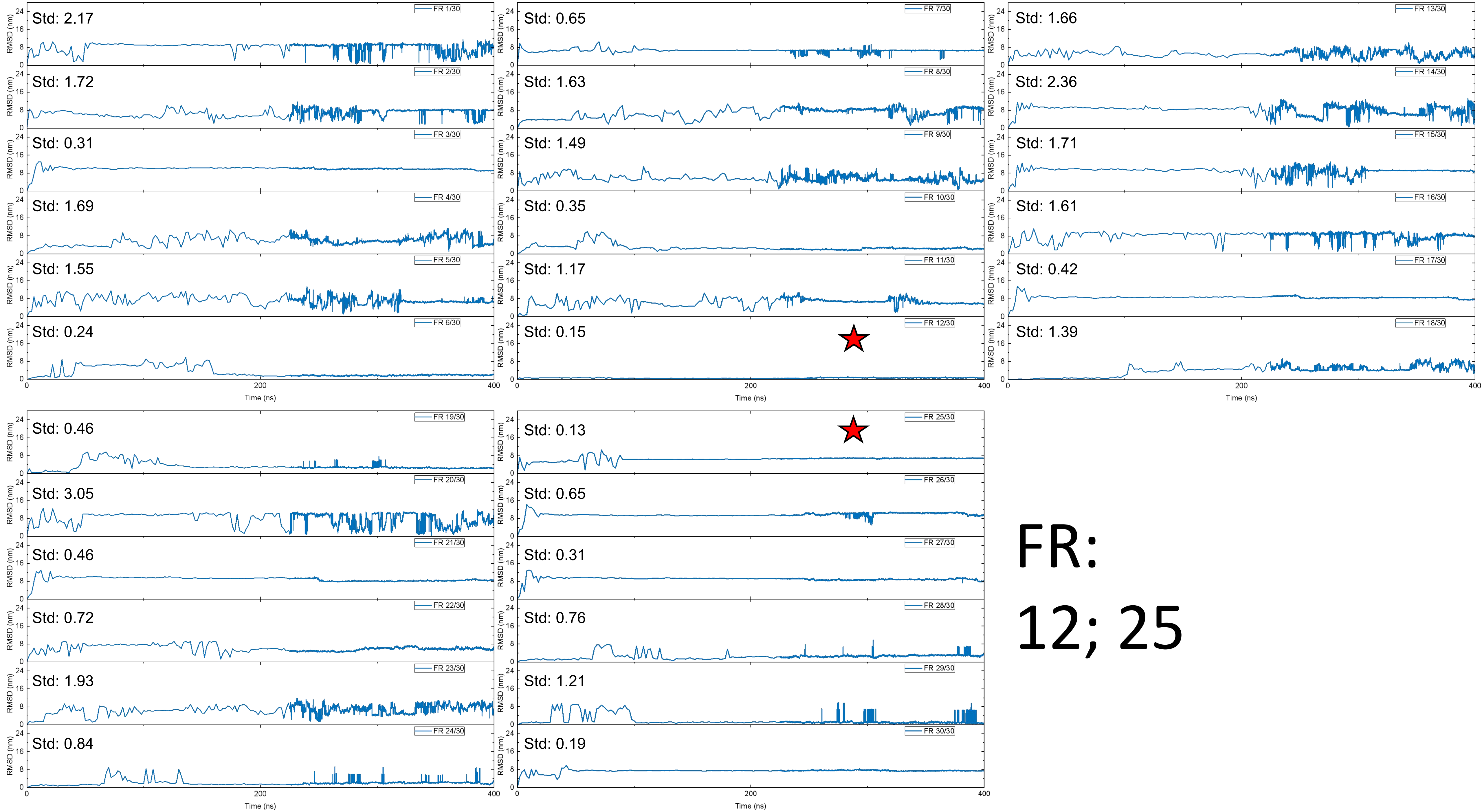

FR 1/30 Std: 2.17
FR 2/30 Std: 1.72
FR 3/30 Std: 0.31
FR 4/30 Std: 1.69
FR 5/30 Std: 1.55
FR 6/30 Std: 0.24
FR 7/30 Std: 0.65
FR 8/30 Std: 1.63
FR 9/30 Std: 1.49
FR 10/30 Std: 0.35
FR 11/30 Std: 1.17
FR 12/30 Std: 0.15
FR 13/30 Std: 1.66
FR 14/30 Std: 2.36
FR 15/30 Std: 1.71
FR 16/30 Std: 1.61
FR 17/30 Std: 0.42
FR 18/30 Std: 1.39
FR 19/30 Std: 0.46
FR 20/30 Std: 3.05
FR 21/30 Std: 0.46
FR 22/30 Std: 0.72
FR 23/30 Std: 1.93
FR 24/30 Std: 0.84
FR 25/30 Std: 0.13
FR 26/30 Std: 0.65
FR 27/30 Std: 0.31
FR 28/30 Std: 0.76
FR 29/30 Std: 1.21
FR 30/30 Std: 0.19
RMSD (nm)
Time (ns)
FR:
12; 25

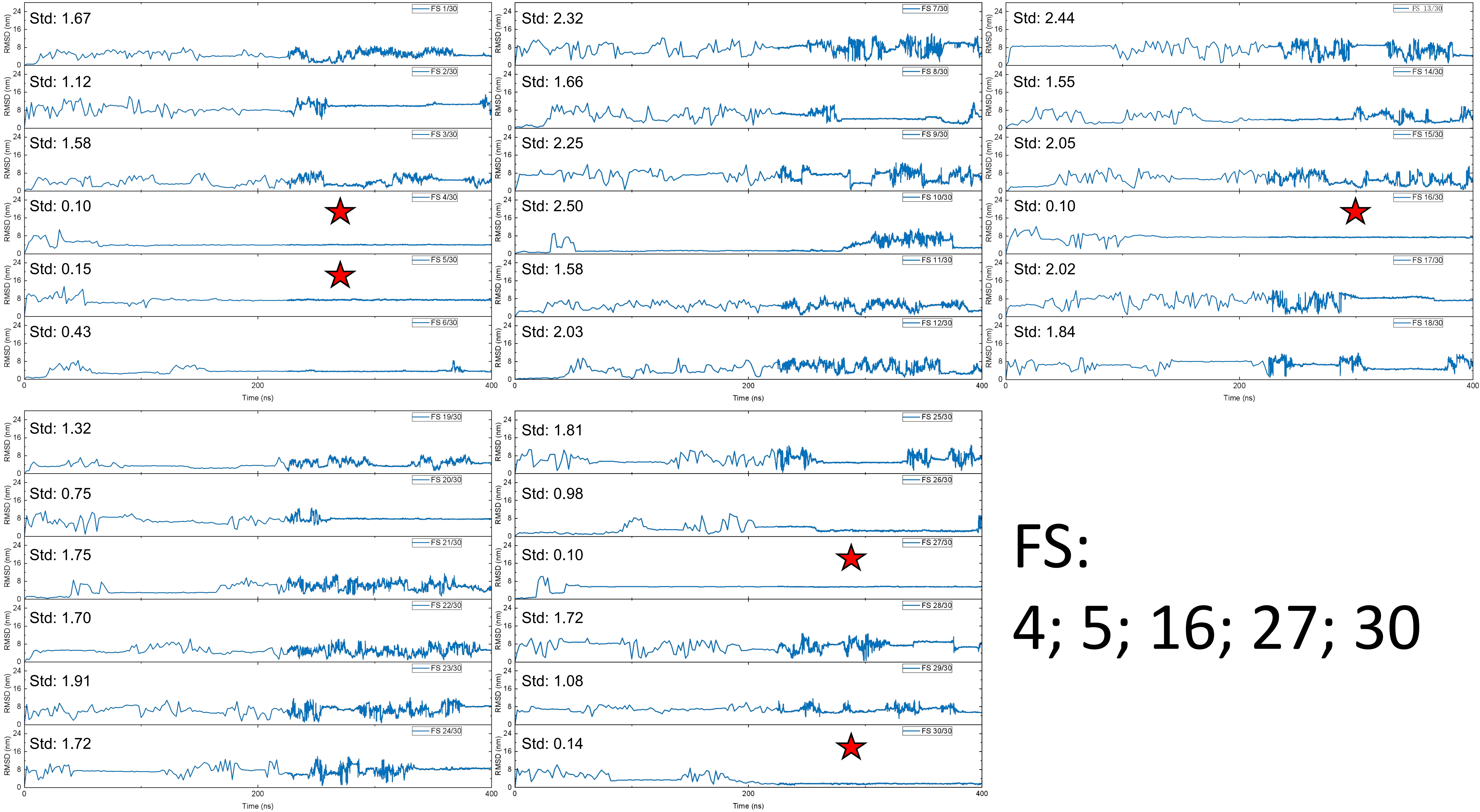

Std: 1.67
FS 1/30
Std: 1.12
FS 2/30
Std: 1.58
FS 3/30
Std: 0.10
FS 4/30
Std: 0.15
FS 5/30
Std: 0.43
FS 6/30
Std: 2.32
FS 7/30
Std: 1.66
FS 8/30
Std: 2.25
FS 9/30
Std: 2.50
FS 10/30
Std: 1.58
FS 11/30
Std: 2.03
FS 12/30
Std: 2.44
FS 13/30
Std: 1.55
FS 14/30
Std: 2.05
FS 15/30
Std: 0.10
FS 16/30
Std: 2.02
FS 17/30
Std: 1.84
FS 18/30
Std: 1.32
FS 19/30
Std: 0.75
FS 20/30
Std: 1.75
FS 21/30
Std: 1.70
FS 22/30
Std: 1.91
FS 23/30
Std: 1.72
FS 24/30
Std: 1.81
FS 25/30
Std: 0.98
FS 26/30
Std: 0.10
FS 27/30
Std: 1.72
FS 28/30
Std: 1.08
FS 29/30
Std: 0.14
FS 30/30
RMSD (nm)
Time (ns)
FS:
4; 5; 16; 27; 30

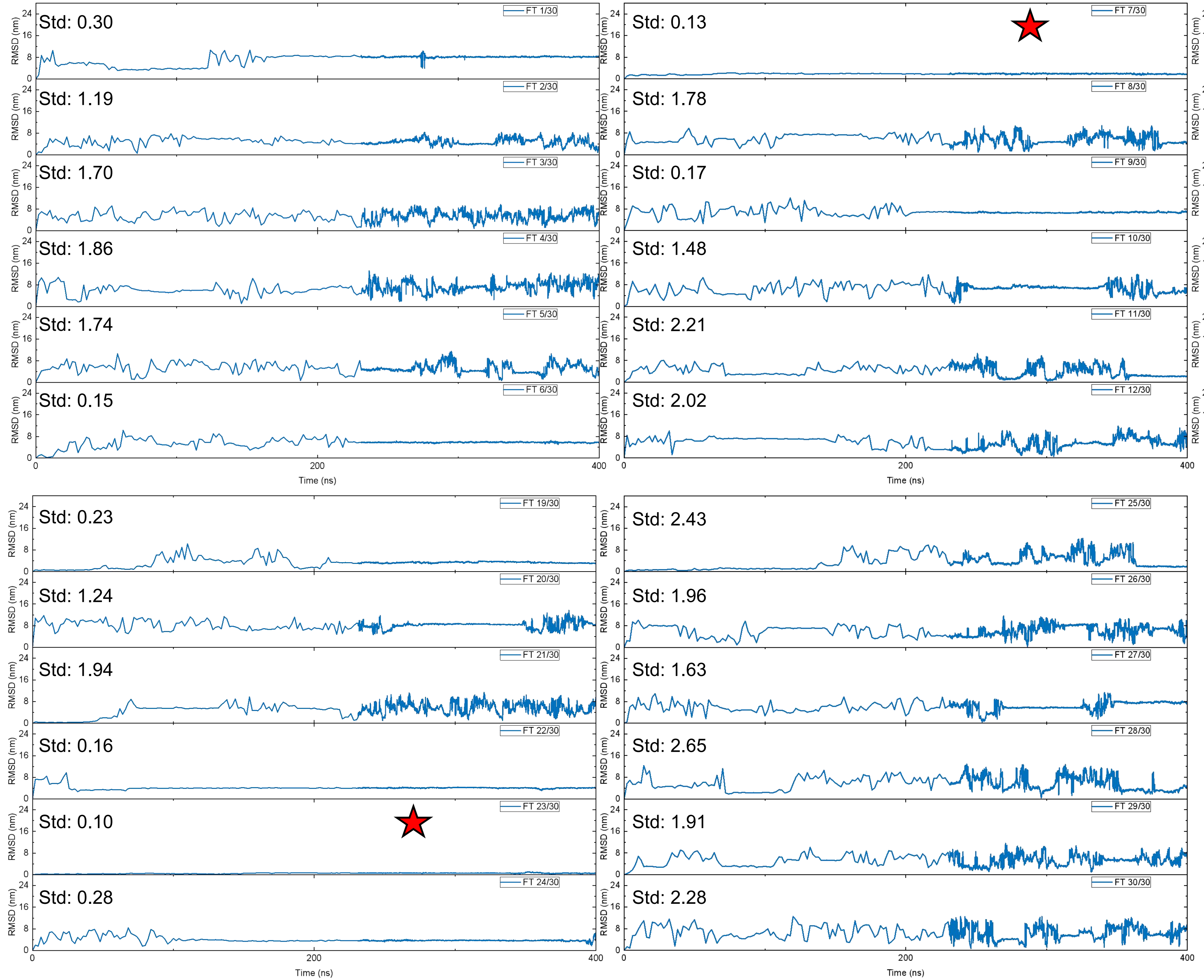

FT 1/30
Std: 0.30
FT 2/30
Std: 1.19
FT 3/30
Std: 1.70
FT 4/30
Std: 1.86
FT 5/30
Std: 1.74
FT 6/30
Std: 0.15
FT 7/30
Std: 0.13
FT 8/30
Std: 1.78
FT 9/30
Std: 0.17
FT 10/30
Std: 1.48
FT 11/30
Std: 2.21
FT 12/30
Std: 2.02
FT 13/30
Std: 1.56
FT 14/30
Std: 0.18
FT 15/30
Std: 2.04
FT 16/30
Std: 2.54
FT 17/30
Std: 1.71
FT 18/30
Std: 0.24
FT 19/30
Std: 0.23
FT 20/30
Std: 1.24
FT 21/30
Std: 1.94
FT 22/30
Std: 0.16
FT 23/30
Std: 0.10
FT 24/30
Std: 0.28
FT 25/30
Std: 2.43
FT 26/30
Std: 1.96
FT 27/30
Std: 1.63
FT 28/30
Std: 2.65
FT 29/30
Std: 1.91
FT 30/30
Std: 2.28
RMSD (nm)
Time (ns)


FT:

7; 23

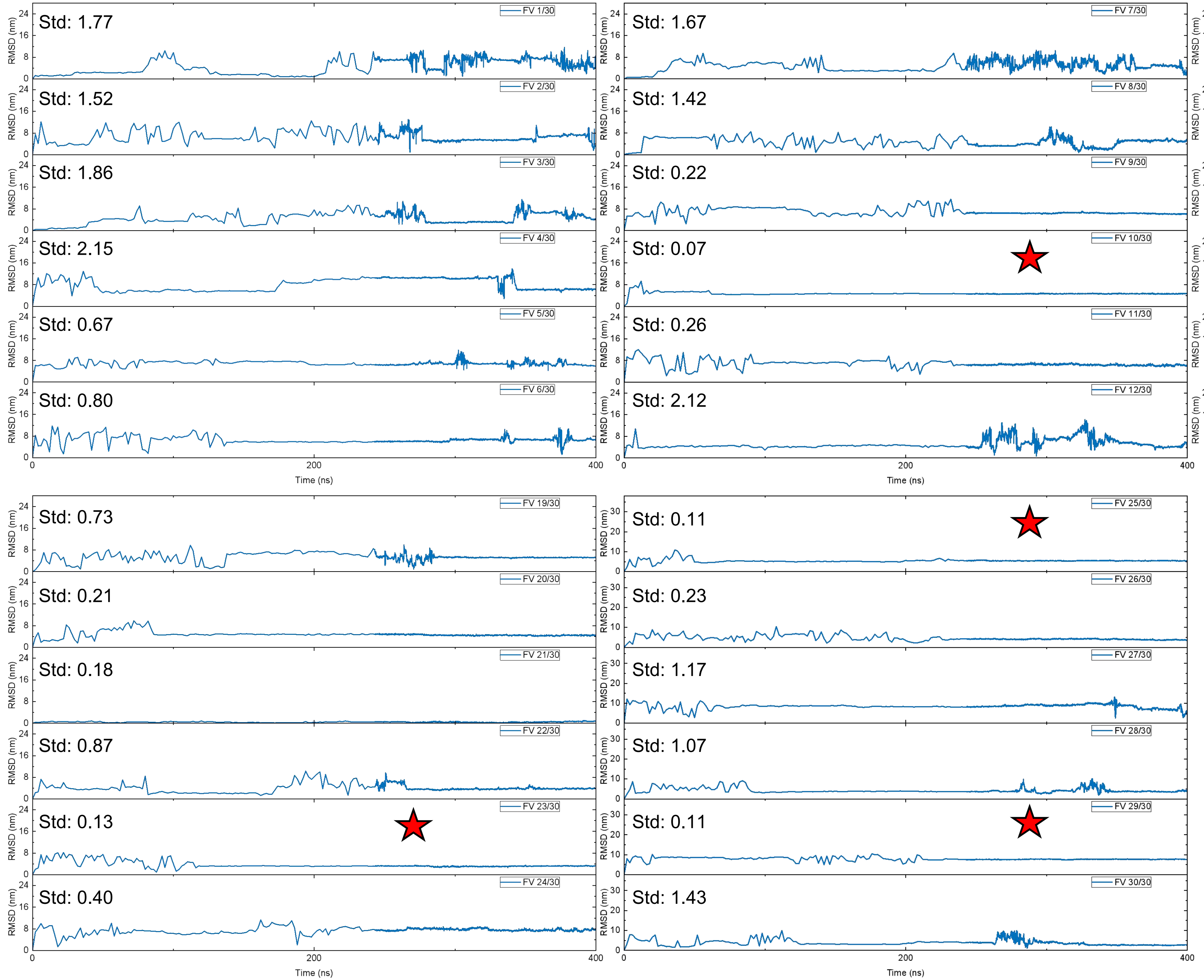


FV:

10; 15; 23; 25;

29

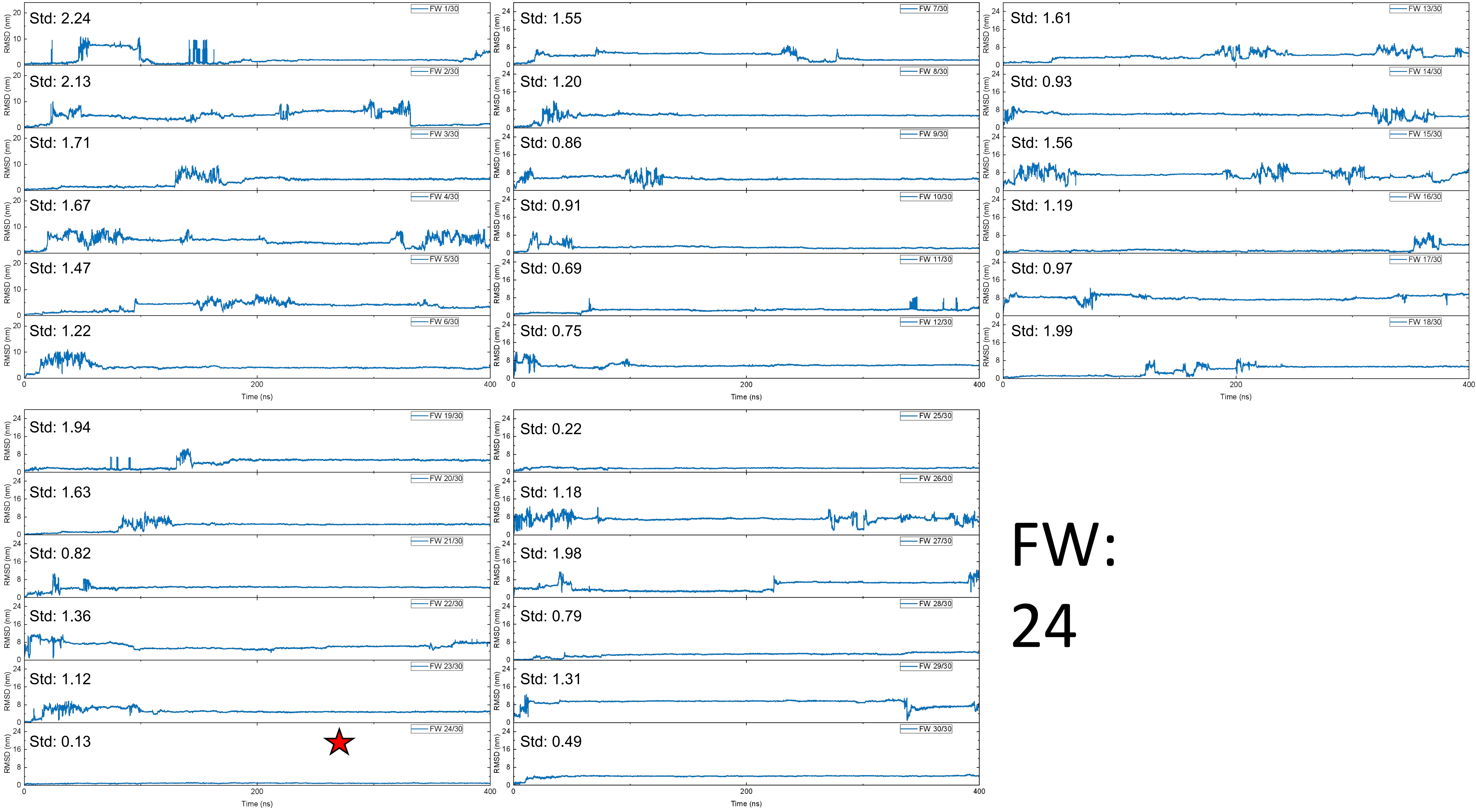
Std: 2.24
FW 1/30
Std: 2.13
FW 2/30
Std: 1.71
FW 3/30
Std: 1.67
FW 4/30
Std: 1.47
FW 5/30
Std: 1.22
FW 6/30
Std: 1.55
FW 7/30
Std: 1.20
FW 8/30
Std: 0.86
FW 9/30
Std: 0.91
FW 10/30
Std: 0.69
FW 11/30
Std: 0.75
FW 12/30
Std: 1.61
FW 13/30
Std: 0.93
FW 14/30
Std: 1.56
FW 15/30
Std: 1.19
FW 16/30
Std: 0.97
FW 17/30
Std: 1.99
FW 18/30
Std: 1.94
FW 19/30
Std: 1.63
FW 20/30
Std: 0.82
FW 21/30
Std: 1.36
FW 22/30
Std: 1.12
FW 23/30
Std: 0.13
FW 24/30
Std: 0.22
FW 25/30
Std: 1.18
FW 26/30
Std: 1.98
FW 27/30
Std: 0.79
FW 28/30
Std: 1.31
FW 29/30
Std: 0.49
FW 30/30
RMSD (nm)
Time (ns)
FW:
24

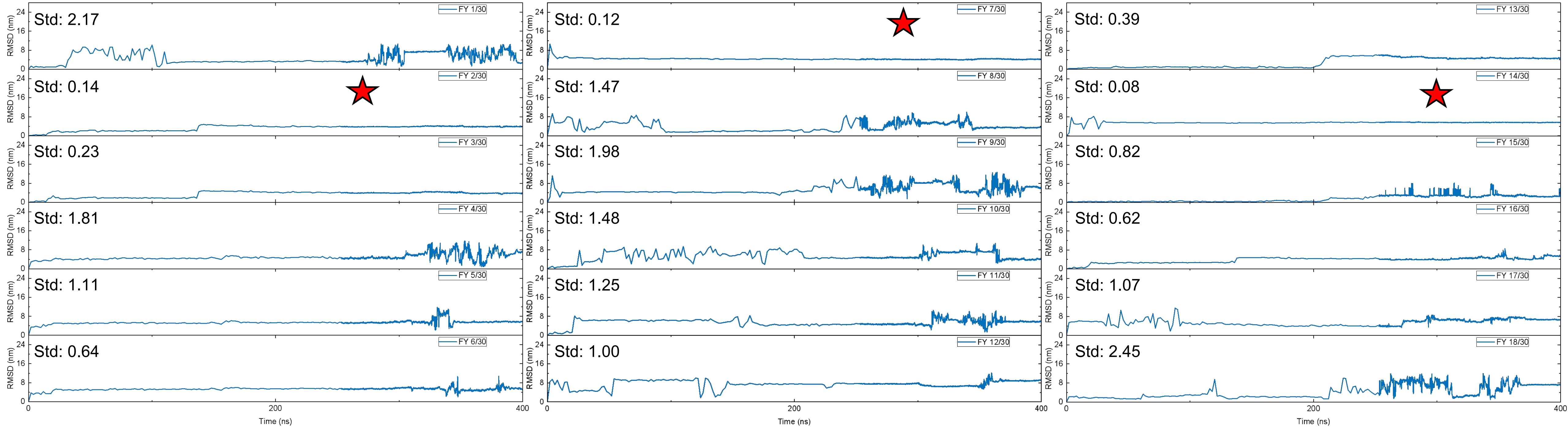


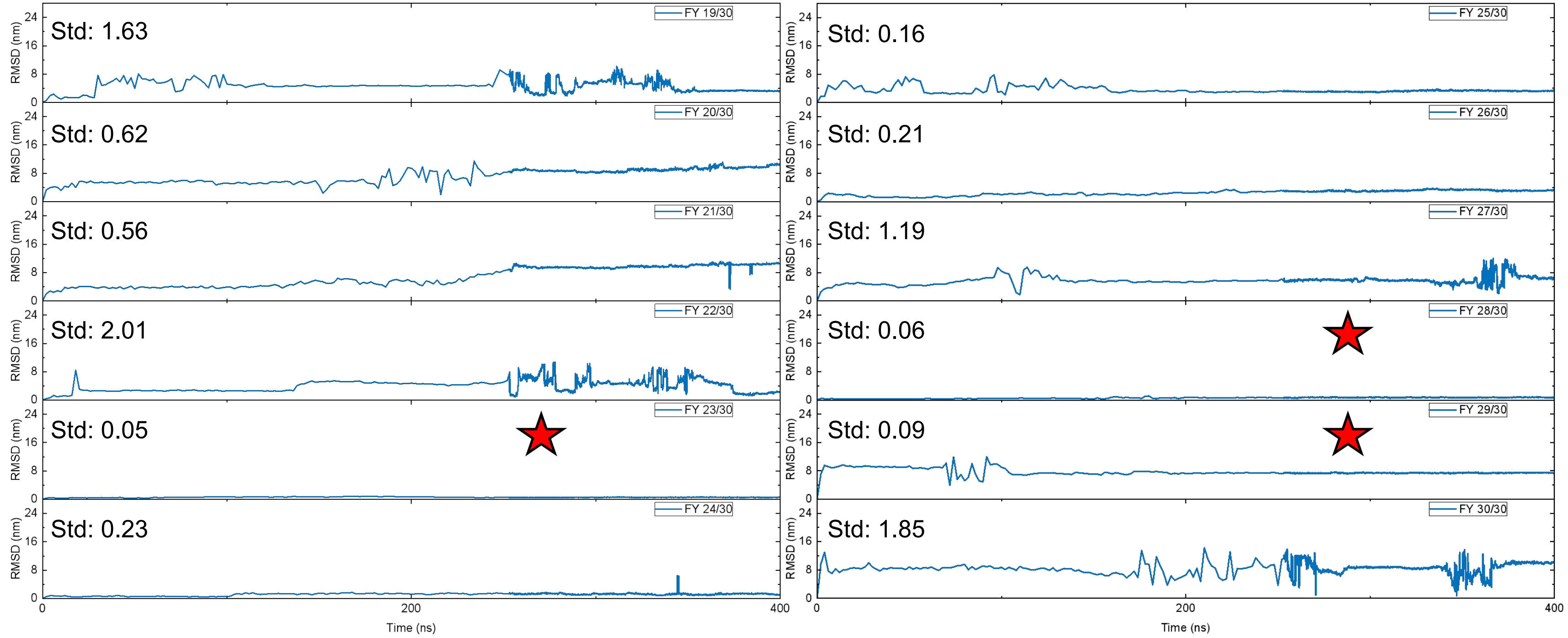


FY:

2; 7; 14; 23; 28; 29
